# Scavenger receptor endocytosis controls apical membrane morphogenesis in the *Drosophila* airways

**DOI:** 10.1101/2022.12.02.518820

**Authors:** Ana S. Pinheiro, Vasilios Tsarouhas, Kirsten Senti, Badrul Arefin, Christos Samakovlis

**Author notes:** Correspondence to C.S. or V.T.

## Abstract

The acquisition of distinct branch sizes and shapes is a central aspect in tubular organ morphogenesis and function. In the *Drosophila* airway tree, the interplay of apical ECM components with the underlying membrane and cytoskeleton controls tube elongation, but the link between ECM composition with apical membrane morphogenesis and tube size regulation is elusive. Here, we characterized Emp (epithelial membrane protein), a *Drosophila* CD36-homologue belonging to the scavenger receptor class B protein-family. *emp* mutant embryos fail to internalize the luminal chitin deacetylases Serp and Verm at the final stages of airway maturation and die at hatching with liquid filled airways. Emp localizes in apical epithelial membranes and shows cargo selectivity for LDLr-domain containing proteins. *emp* mutants also display over elongated tracheal tubes with increased levels of the apical proteins Crb, DE-cad and phosphorylated Src (p-Src). We show that Emp associates and organizes the βH-Spectrin cytoskeleton and is itself confined by apical F-actin bundles. Overexpression or loss of its cargo protein Serp lead to abnormal apical accumulations of Emp and perturbations in p-Src levels. We propose that during morphogenesis, Emp senses and responds to luminal cargo levels by initiating apical membrane endocytosis along the longitudinal tube axis and thereby restricts airway elongation.

## Introduction

The tube shapes in transporting organs like kidney, lung and vascular system are precisely controlled to ensure optimal fluid flow and thereby, function. Failure in normal tube size acquisition leads to cystic, stenotic or winding tubes. The *Drosophila* respiratory network, the trachea, provides a well-characterized system for the genetic dissection of tubular organ maturation. Like mammalian lungs, the trachea undergoes a precisely timed series of maturation events to convert the nascent branches into functional airways. First, a transient secretion burst of luminal proteins (Tsarouhas *et al*., 2007; Jayaram *et al*., 2008; Förster, Armbruster and Luschnig, 2010) and initiates tube expansion. Luminal proteins assemble into a chitinous central rod and into an apical ECM (taenidia) lining the apical membrane. After 10 hours, luminal material becomes rapidly cleared from the tubes by massive endocytosis involving several endocytic pathways (Tsarouhas *et al*., 2007). Finally, a liquid clearance pulse converts the tubes into functional airways (Tsarouhas *et al*., 2007). Genetic studies suggested an instructive role of luminal chitin and proteins in tube growth coordination and termination. Mutations affecting chitin biosynthesis (kkv) or matrix assembly (knk, gasp) show irregular tube shapes, diametric expansion and tube maturation defects (Moussian *et al*., 2006; Tiklová, Tsarouhas and Samakovlis, 2013; Öztürk-Çolak *et al*., 2016). Tube elongation is continuous during tracheal development and its termination requires chitin biosynthesis and the secreted chitin deacetylases vermiform (Verm) and serpentine (Serp) (Beitel and Kransnow, 2000; Luschnig *et al*., 2006; Wang *et al*., 2006). These luminal proteins presumably modify the structure and physical properties of the extra-cellular matrix (ECM) and thereby restrict tube elongation. In addition to the luminal matrix pathway, components involved in the assembly of basolateral septate junctions (SJs) also restrict tube elongation, through the regulation of the subcellular localization of Crumbs, a transmembrane protein that promotes expansion of the tracheal cell apical surface and tube elongation (Laprise *et al*., 2010). More recently, the conserved non-receptor tyrosine kinase Src42 was found to promote axial elongation by controlling the apical cytoskeleton and apical cell junctions (Förster and Luschnig, 2012; Nelson *et al*., 2012; Olivares-Castiñeira and Llimargas, 2018). Additionally, Yki and several components of the Hippo pathway control tube elongation, along with transcription factors like Blimp-1 and Grh (Hemphälä *et al*., 2003; Robbins, Gbur and Beitel, 2014; Öztürk-çolak *et al*., 2018; McSharry and Beitel, 2019; Skouloudaki *et al*., 2019). An appealing model suggests that the interaction between apical membrane and ECM elasticity may influence apical cytoskeletal organization and thereby control tube shapes (Dong, Hannezo and Hayashi, 2014). Although ECM integrity and the apical cytoskeleton appear crucial in tube length regulation, it is unknown how ECM signals are perceived by the airway cells to regulate their shapes during tube maturation.

Scavenger receptors comprise a superfamily of cell surface membrane proteins that bind and internalize modified lipoproteins and various other types of ligands. CD36 (cluster of differentiation 36) belongs to class B scavenger receptor family, which includes scavenger receptor B1 (SRB1) and lysosomal integral membrane protein 2 (LIMP2). CD36 is expressed on the surface of many cell types including epithelial, endothelial cells, and macrophages. Disruption of CD36 function in mice can lead to inflammation, atherosclerosis, metabolic disorders, tumor growth and metastasis (Chen *et al*., 2008; Pascual *et al*., 2017; Wang *et al*., 2020). CD36 has several cargoes, including long-chain fatty acids, oxidized LDL (ox-LDL), oxidized phospholipids and thrombospondin-1 (TSP-1) (Githaka *et al*., 2016; Yang *et al*., 2017; Deng *et al*., 2022). In vitro imaging studies of macrophages and endothelial cells propose that CD36 clustering at the cell surface upon engagement of multivalent ligands and in conjunction with the cortical cytoskeleton triggers signal transduction and receptor-ligand complex endocytosis (Jaqaman *et al*., 2011; Githaka *et al*., 2016). The activity of several signaling effectors, including the Src family kinases, Fyn, Yes (Thorne *et al*., 2006; Zani *et al*., 2015) and the mitogen-activated kinases, Jun-kinase (JNK) 1 and 2 (Rahaman *et al*., 2006) can be regulated by CD36. The *Drosophila* genome includes a family of 14 CD36-like genes, with distinct tissue-specific expression patterns. The genetic analysis of a few members in this class B scavenger receptor family implicated them in phagocytosis, immune responses and photoreceptor function (Philips, Rubin and Perrimon, 2005; Stuart *et al*., 2005; Voolstra *et al*., 2006). The *Drosophila* Emp (Epithelial membrane protein) shows the highest similarity with CD36 and is selectively expressed in embryonic epithelial tissues (Hart, Klein and Wilcox, 1993).

Here, we show that Emp is a selective receptor for internalization, endosomal targeting and tracheal luminal clearance of proteins with LDLr domains. *emp* mutants display over elongated tracheal tubes with increased levels of junctional Crb, DE-cad and phospho-Src. Reduction of Src42A in *emp* mutants, rescues the tube elongation phenotype indicating that Emp modulates junctional *p*-Src42A levels to control apical membrane expansion and tube length. The organization of the beta-heavy spectrin (*β*H-Spectrin) cytoskeletal network is compromised in *emp* mutants. Emp binds to *β*H-Spectrin suggesting that it provides a direct link between ECM, apical membrane and cytoskeleton during tube maturation process. Re-expression of human CD36 in *emp* mutants can ameliorate the mutant tube phenotypes suggesting conserved functions of Emp.

## Results

### Emp is a selective scavenger receptor required for tube elongation and luminal protein clearance

*emp* (or CG2727 in Flybase) encodes a class-B scavenger receptor expressed in embryonic ectodermal epithelial tissues including the tracheal system (Hart, Klein and Wilcox, 1993). To elucidate the developmental functions of *emp* in the airways, we generated a deletion mutant (*emp^e3d1^,* referred as *emp* mutant hereafter) using the FLP/FRT recombinase system (Parks *et al*., 2004) (Figure 1-figure supplement 1A). PCR mapping of genomic DNA identified a 4.8 kb deletion encompassing exon 2 in the *CG2727* locus. Both immunofluorescence and Western blots using a polyclonal antiserum against recombinant Emp (see Methods) (Figure 1-figure supplement 1B), failed to detect Emp protein in *emp* mutants (Figure 1-figure supplement 1C-D). Similarly, quantitative RT-PCR of RNA extracted from late embryos showed a strong reduction of *emp* RNA in the mutants (Figure 1-figure supplement 1E). *emp* homozygous or *Df(2R)BSC608*/*emp* mutants were embryonic lethal with few escapers surviving to 1^st^ instar larvae. This embryonic lethality could be rescued by the re-expression of a transgenic *emp* construct using the ectodermal driver *69BGal4.* This suggests that the deletion generates a strong loss-of function mutation in *emp* and does not affect any neighboring genes required for embryo viability. To examine a potential role of Emp in airway maturation, we visualized the tracheal tubes during embryonic development in *wild-type* and *emp* mutants. *emp* embryos at stage 16 showed a 30% over-elongation of the dorsal trunk (DT) compared to the *wild-type* (Figure 1A), but showed no defects in tube diameter (Figure 1-figure supplement 1F). *emp* mutants also failed to fill their airways with gas at hatching (Figure 1-figure supplement 1I, J). Both phenotypes could be rescued by re-expression of *emp* in tracheal cells of the *emp* mutants (Figure 1A, 1B, Figure 1-figure supplement 1 Ic, J). We conclude that Emp is required for normal tube elongation and gas-filling during embryonic development. The survival of *emp* mutants overexpressing *emp* in the airways was limited to larval stages, suggesting that the tracheal-specific re-expression of *emp* is not sufficient for larval or adult survival. This suggests additional roles for Emp in other tissues. The human homologue of Emp, CD36 shares the overall protein architecture and 30% of amino acid identity with Emp (Figure 1-figure supplement 1 G, H). We generated a transgenic line expressing the coding sequence of human CD36 and drove its expression in fly airways to test if Emp and CD36 have conserved functions. We found that both the tracheal length and gas-filling defects in *emp* mutants were partially reversed by tracheal CD36 overexpression, arguing for a conserved function of CD36 (Figure 1A, 1B, Figure 1-figure supplement 1Id, J).

**Figure 1.**
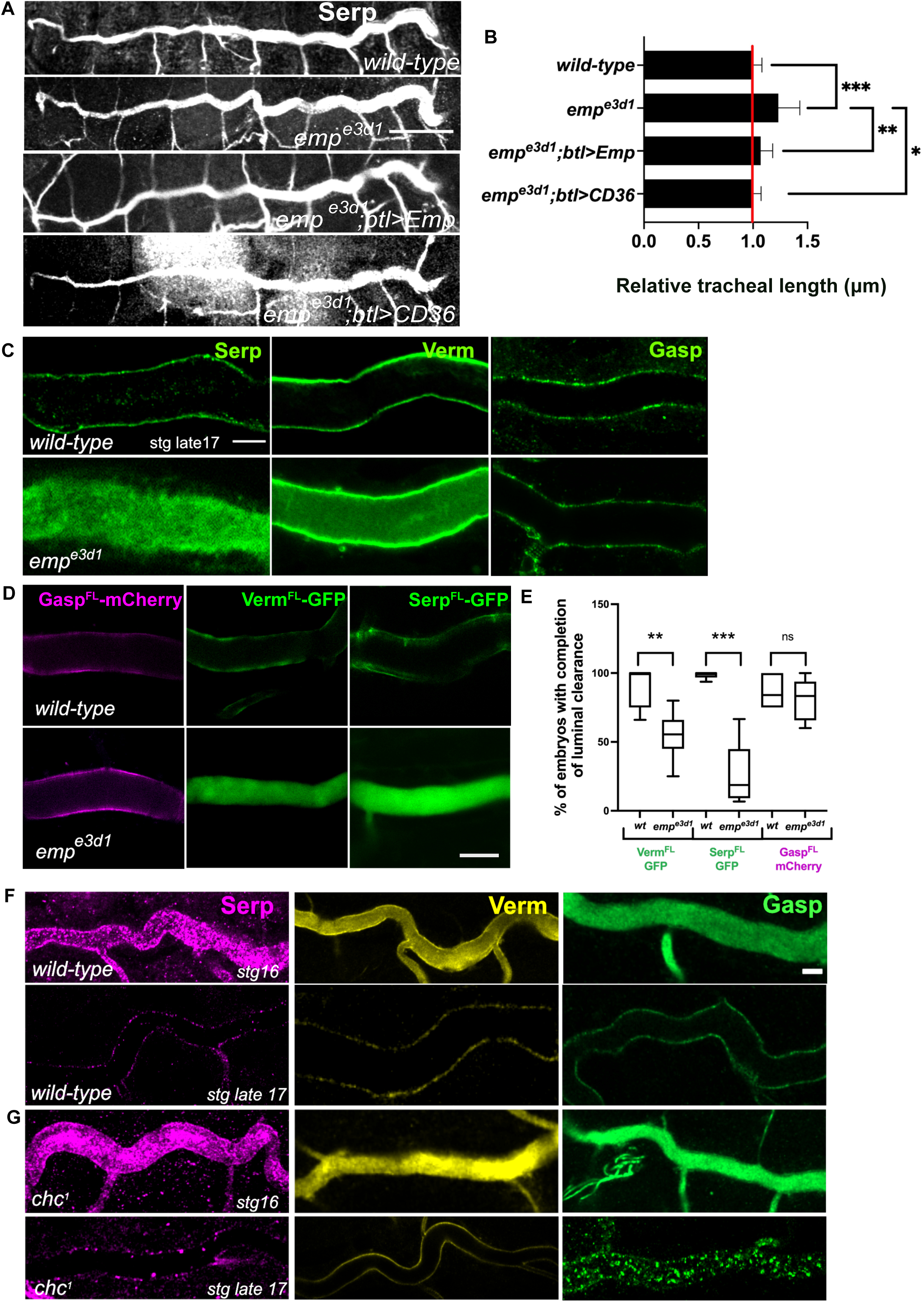
*emp^e3d1^* mutants show over-elongation of the tracheal tube and severe luminal clearance defect of Serp. (A) Images showing the DT of *wild-type*, *emp^e3d1^*, *emp^e3d1^;btl>Emp*, *emp^e3d1^;btl>CD36* embryos stained for the luminal marker Serp. (B) Graph showing the tracheal DT length in *wild-type* embryos (n=16), *emp^e3d1^* mutants (n=17), *emp^e3d1^;btl>Emp* (n=19) and *emp^e3d1^;btl>CD36 (n=15)* embryos. (C) Confocal images showing the DT of *wild-type* and *emp^e3d1^* mutant at late stage 17 embryos, stained for the endogenous luminal proteins Serp, Verm and Gasp. (D) Confocal images showing the DT of live *btl>Gasp^FL^-mCherry* (magenta), *btl>Verm^FL^-GFP* and *btl>Serp^FL^-GFP* (green) in *wild-type*, and *emp^e3d1^* mutant at 20.0 h AEL. (E) Plots showing the percentage of embryos with completion of luminal clearance in *btl>Verm^FL^-GFP* (green, n=59), *emp^e3d1^*;*btl>Verm^FL^-GFP* (green, n=37), *btl>Serp^FL^-GFP (green, n=58), emp^e3d1^;btl>Serp^FL^-GFP* (green, n=15), *btl>Gasp^FL^-mCherry* (magenta, n=28) and *emp^e3d1^*;*btl>Gasp^FL^-mCherry* (magenta, n=18). (F) Confocal images showing the tracheal DT of *wild-type* and *chc^1^* mutant embryos, stained for the endogenous luminal markers Serp (magenta), Verm (yellow) and Gasp (green) before and after luminal clearance, stage 16 and late stage 17. Error bars denote s.e.m., *p < 0,05* p < 0,005\*\** and *p < 0,0005\*\*\** (unpaired two tailed t tests). Scale bars, 50μm, 10μm and 5μm for images (A), (C, D) and (F) respectively.

To establish whether the failure of tracheal gas-filling originates from a defect in luminal protein clearance, we stained *wild-type* and *emp* mutants for the luminal proteins Serp, Verm and Gasp. These secretory proteins were internalized from the lumen by late stage 17 in *wild-type* embryos. Serp and Verm, but not Gasp were selectively retained in the *emp* dorsal trunk (DT) airways (Figure 1C) suggesting a role of Emp in the internalization of a subset of luminal proteins. To confirm the protein clearance phenotypes, we generated *emp* mutants carrying *btl>Serp-GFP* or *btl>Verm-GFP* or *btl>Gasp-mCherry* transgenes and analyzed them by live-imaging from 17h to 21h AEL (Figure 1D, E). The three reporters were normally secreted into the lumen of both *emp* mutants and *wild-type* embryos and at 19 hours they were cleared from the tubes of *wild-type* embryos. In *emp* mutants *Gasp-mCherry* was also cleared from the lumen but the Serp-GFP and Verm-GFP reporters remained inside the tube. This suggests that Emp acts as a selective endocytosis receptor during luminal protein clearance. The unperturbed clearance of Gasp-Cherry suggested differential requirements for the internalization of luminal proteins. We stained *clathrin* (*chc^1^)* mutants for Serp, Verm and Gasp and analyzed them at late embryonic stages, when luminal clearance is completed in *wild-type* embryos. We found that Serp and Verm were cleared, whereas Gasp clearance was selectively impaired in *chc^1^* mutants (Figure 1F). This indicates that Emp is involved in a selective, clathrin-independent endocytosis pathway internalizing Verm and Serp. Gasp endocytosis is clathrin-dependent and presumably involves an unidentified surface receptor. These genetic experiments suggest that Emp controls tracheal tube elongation and luminal clearance of chitin deacetylases presumably by mediating their endocytosis.

### Dynamic subcellular localization of Emp during tracheal development

To further elucidate Emp functions, we generated an antibody against its extracellular domain (Figure 1-figure supplement 1B) and determined its subcellular localization by co-staining for the previously characterized apical membrane protein Crb and markers of adherens (DE-cad, pY) and septate epithelial junctions (Dlg, Cora). During tracheal branch elongation (stage 14 to 16), *wild-type* embryos showed an Emp enrichment in epithelial apical membranes and in subapical cytoplasmic puncta (Figure 2A). Similar to the tracheal cells, we detected apical Emp localization, initially diffuse in dots in subapical regions and progressively more defined at the apical cortical region in the epidermis and hindgut of stage 15 to 16 *wild-type* embryos (Figure 2-figure supplement 1A-B). During late stage 16 - early stage 17, Emp localization became predominantly restricted in the junctional subapical region of epithelial cells, where it colocalized with Crb, DEcad-GFP, and Phospho-Src (Figure 2A, Figure 2-figure supplement C-D). The Emp signal showed only weak colocalization with the septate junction markers Coracle, Mtf, Dlg and with the lateral cytoskeleton marked by *α*-Spectrin (Figure 2-figure supplement 1 A-C). The massive uptake of luminal material correlates with the disassembly of apical actin bundles running along the transverse tube axis. The diaphanous-like formin, DAAM and the type III receptor tyrosine phosphatases, Ptp4E and Ptp10D, control the organization of F-actin bundles running along the perpendicular tube axis in the *Drosophila* airways (Matusek *et al*., 2006; Tsarouhas *et al*., 2019). Mutations in *Ptp10D4E* (*Ptp10D* and *Ptp4E*) or expression of a dominant negative form of DAAM (*btl*>C-DAAM) disrupt the transverse actin bundle arrays and prematurely initiate luminal protein clearance. Similarly, Latrunculin A (Lat-A) injection in embryos destroys the actin bundles and leads to luminal protein uptake. These experiments had suggested that the transverse F-actin bundles restrict the endocytic uptake of luminal cargoes (Tsarouhas *et al*., 2019). We tested whether the dynamic localization of Emp may be altered in *Ptp10D4E* mutants and in embryos overexpressing the dominant negative C-DAAM construct in the airways. Embryos of both genotypes showed premature translocation of Emp to the airway cell junctions compared to *wild-type* (Figure 2B, C). This suggests, that similar to luminal protein uptake, the relocation of Emp to the apical junctional region can be induced by the premature disassembly of the actin bundles. Additionally, we analyzed the localization and intensity of Emp and Crb protein stainings along the apical membrane in *wild-type* and *btl*>C-DAAM embryos. DAAM inactivation increased the intensity of apical Emp punctuate accumulations compared to *wild-type* (Figure 2E, F), further arguing that the transverse actin bundles restrict Emp localization at the apical membrane.

**Figure 2.**
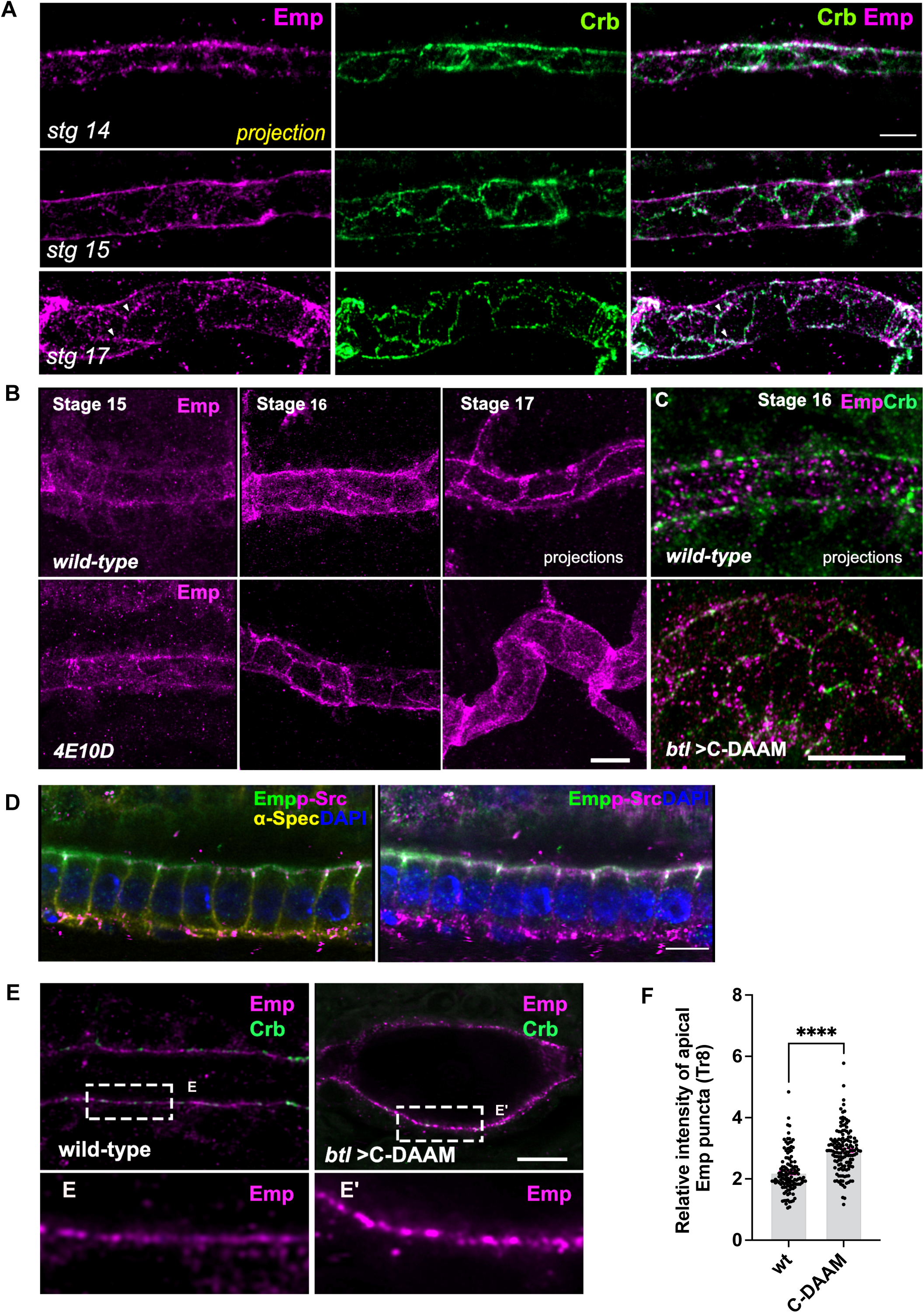
Dynamic apical distribution of Emp during tube maturation. (A) Confocal images showing dorsal trunk (DT) projections, from stage 14, 15 and 17 embryos, stained for Emp and Crb (Crumbs). (B) DT projections, from stage 15, 16 and 17 of *wild-type* and *Ptp4E10D* (4E10D) stained for Emp. (C) Confocal images of the DT in *wild-type* and *btl>C-DAAM* stained for Emp and Crb. (D) Confocal images of embryonic tracheal cells stained for Emp, p-Src, *α*-Spectrin (*α*-Spec) and DAPI showing the subcellular localization of Emp in stage 17. (E) Confocal images showing the tracheal DT of *wild-type* and *btl>C-DAAM* embryos stained for Emp and Crb. Insets shows the magnification of (E), indicated by the white rectangles. (F) Bar plot showing the relative intensity of apical Emp puncta at DT (Tr8) in *wild-type* (*n* = 168 puncta, 5 embryos) and *btl>C-DAAM* (*n* = 237 puncta, 6 embryos). Statistically significant shown in *p* - values p < 0,0005***(unpaired two tailed t tests). Scale bars, 5μm (A, B, D, E) and 10 μm (C).

To test if a subset of the apical Emp puncta may correspond to endocytic vesicles, we analyzed the localization of Emp relatively to several YFP-tagged Rab GTPases (YRab), expressed at endogenous levels. Co-staining for Emp and GFP (Dunst *et al*., 2015) showed an overlap with YRab5 (early endosomes) and YRab7 (late endosomes) with the Emp positive cytoplasmic puncta. We also detected weaker colocalization with YRab11 (recycling endosomes), (Figure 3A). Live-imaging of embryos expressing *btl>Emp-GFP* in the time interval of luminal protein clearance (early stage 17) showed an increase of Emp intracellular puncta compared to stage15 or late stage 17 embryos (Figure 2-figure supplement 1D). Overall, these experiments suggest that the localization of Emp in the apical membrane and endocytic vesicles is dynamic and influenced by actin bundle integrity. The timing of the final, steady-state accumulation of Emp is controlled by PTP signaling.

**Figure 3.**
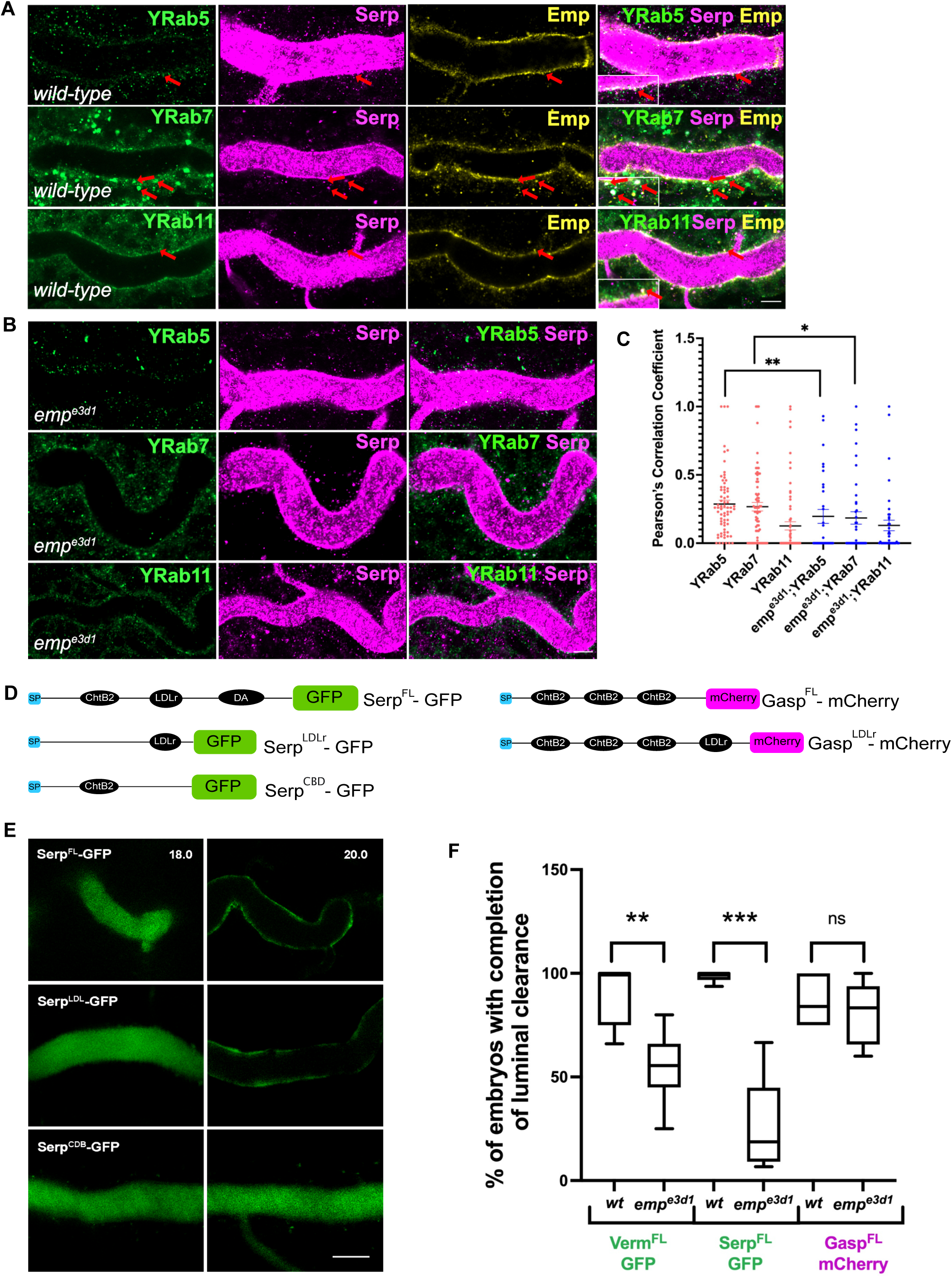
The endosomal localization of Serp is strongly reduced in *emp^e3d1^* mutants. (A) Confocal images of tracheal DT of *wild-type* embryos expressing endogenous tagged YFP-Rab (knock-in) proteins, YRab5, YRab7 and YRab11 stained with Emp, Serp and GFP. Insets show zoomed cross section views of the DT (*y-z* plane) of Emp co-localization with YRab5, YRab7 and YRab11, as indicated with re d arrowheads. (B) Confocal images showing the tracheal DT, stained for Serp and endogenous YRab5, YRab7 and YRab11 in *emp^e3d1^* mutant. (C) Scatter plots representing the co-localization between the YRabs and Serp in *wild-type* and in *emp^3ed1^* mutants. (D) Schematic representation of Serp and Gasp domain organization. The following abbreviations are used: SP, signal peptide (blue); LDLr, low density lipoprotein receptor (black); ChtB, Chitin binding domain (black); GFP (green); Cht BD2, chitin binding domain (black); and mCherry (magenta). *btl>Serp^FL^-GFP* represents the full length of Serp, *btl>Serp^LDLr^-GFP* represents the LDLr domain of Serp, *btl>Serp^CBD^-GFP* expresses ChtB domain of Serp, *btl>Gasp^FL^-mCherry* represents the full-length Gasp protein *and btl>Gasp^LDLr^-mCherry* represents the full-length Gasp protein with addition of the LDLr domain. (E) Confocal images showing the DT of live *btl>Serp^FL^-GFP, btl>Serp^LDL^-GFP, btl>Serp^CBD^-GFP,* (green) embryos before (18.0 h AEL) and after (20.0 h AEL) luminal protein clearance. *btl>Serp^CBD^-GFP* embryos show incomplete luminal GFP clearance compared to *btl>Serp^FL^-GFP or btl>Serp^LDLr^-GFP*. (F) Plots showing the percentage of embryos with completion of luminal clearance in *btl>Serp^FL^-GFP* (green, n = 58); *emp^e3d1^*;*btl>Serp^FL^-GFP* (green, n = 15); *btl>Serp^LDLr^-GFP* (green, n = 32); *emp^e3d1^;btl>Serp^LDLr^-GFP* (green, *n* = 2 6); *btl>Serp^CBD^-GFP* (green, *n* = 45); *emp^e3d1^*; *btl>Serp^CBD^-GFP* (green, n = 28); *btl>Gasp^Fl-LDLr^-mCherry* (magenta, *n* = 57); and *emp^e3d1^*;*btl>Gasp^FL-LDLr^-mCherry* (magenta, *n* = 56). The median (horizontal line) is shown in the plots with the data range from 25^th^ to 75^th^ percentile. Error bars denote s.e.m., p < 0,05*, and p < 0,0005*** (unpaired two tailed *t* tests). Scale bars, 5 μm (A, B) and 10 μm (E).

### Serp internalization and endosomal targeting requires Emp

The luminal retention of Serp in *emp* mutants and the partial localization of Emp with endosomal markers led us to examine if Emp mediates the endosomal uptake and trafficking of luminal Serp. We co-stained for endogenous YFP-tagged endosomal markers and Serp in *wild-type* and *emp* mutant embryos. This analysis showed that intracellular Serp puncta co-stained for the early endosomal marker Rab5 (Figure 3A, C red arrows, R=0.29) and late endosomal marker, Rab7 (Figure 3A, C red arrows, R=0,27) in *wild-type* embryos. In the *emp* mutants the colocalization of Serp with both early and late endocytic markers was significantly decreased (Figure 3B), suggesting that Emp mediates Serp internalization and endosomal vesicle targeting. In addition, the number of intracellular Serp puncta were reduced in *emp* mutant embryos compared to *wild-type* (Figure 3-figure supplement 1A), whereas the total number of GFP puncta corresponding to early and late endosomes remained unchanged (Figure 3-figure supplement 1B). Taken together, these results suggest that Emp is a receptor for Serp internalization and endosomal targeting.

To further investigate Emp cargo specificity, we tested the luminal clearance of GFP constructs, tagged with different domains of Serp in *wild-type* embryos and *emp* mutants (Figure 3D). We used GFP constructs fused to either Serp-full-length or to the Serp-LDLr-domain (Low density lipoprotein receptor-domain) or to the Serp-CBD-domain (chitin binding domain) (Luschnig *et al*., 2006; Wang *et al*., 2006) and examined their luminal secretion and clearance. The constructs were expressed and normally secreted into the tracheal tubes of *wild-type emp* mutant embryos. The Serp-LDLr reporter was cleared from the lumen as efficient as the full-length Serp-GFP but the CBD-GFP fusion was retained in the tracheal lumen of 20% of *wild-type* embryos. Interestingly both the Serp-GFP and LDLr-GFP were strongly retained in the lumen of *emp* mutants. These results suggest that the LDLr domain of Serp, targets GFP to Emp-mediated internalization (Figure 3E, F). CBD-GFP clearance failed in 40% of the *emp* mutants suggesting that this cargo is also internalized by alternative receptors. To further test if the addition of the LDLr-domain is sufficient to target an unrelated protein for Emp-mediated uptake, we fused the Serp LDLr-domain to the Gasp-mCherry protein, which does not require *emp* for its luminal clearance. As with the Serp-based constructs we analyzed the clearance of *Gasp^FL^-mCherry* and *Gasp^FL+LDLr^-mCherry* in *wild-type* and *emp* mutant embryos (Figure 3D). Both constructs were normally cleared form the airways of *wild-type* embryos. However, *btl>Gasp^FL+LDLr^-mCherry,* but not *btl>Gasp^FL^-mCherry,* was retained in the airways of *emp* embryos (Figure 3F compare with Figure 1E). These data suggest that the LDLr domain confers cargo specificity for Emp-mediated internalization. Loss of function *verm serp* mutants or overexpression of *serp* leads to over elongation of the tracheal tubes (Luschnig *et al*., 2006; Wang *et al*., 2006). We thus examined the effects of Serp and Verm on the levels and localization of Emp. *btl>Serp-GFP* overexpressing embryos showed increased punctate accumulations of Emp and Crb at the apical cell surface compared to *wild-type* (Figure 4A). Conversely, in *verm^ex245^,serp^ex7^* double mutants, we detected more diffuse punctate cytoplasmic accumulations for both Emp and Crb compared to *wild-type* (Figure 4A, B). To investigate whether the changes in intensity and localization of Emp and Crb puncta upon Serp-GFP overexpression or *verm* and *serp* deletion were due to changes in protein synthesis or stability we performed Western blots of mutant and *wild-type* embryos. The total protein levels of Crb and Emp did not change in the mutants suggesting that the levels of luminal Serp control the punctate accumulation of Emp and Crb at the apical membrane (Figure 4D-F). As expected, overexpression of Emp in the airways (*btl>Emp*) increased the overall levels of cytoplasmic Emp, but did not influence the accumulation of Crb (Figure 4D). Overall, these data suggest that the levels of luminal cargo control the punctate accumulation of Emp and Crb at the apical membrane. Measurements from the tr5 to tr10 showed reduced DT elongation in *emp^e3d1^;btl>Serp-GFP* embryos compared to *btl>Serp-GFP*, suggesting that Serp overexpression control the length of the *Drosophila* airways at least partially through Emp (Figure 4G). Overall, these results suggest that Emp acts as a scavenger receptor for LDLr domain proteins, such as Serp thereby facilitating their internalization through clathrin independent endocytosis. The levels of luminal Serp control the localized apical membrane accumulation of Emp and Crb to calibrate tube elongation.

**Figure 4.**
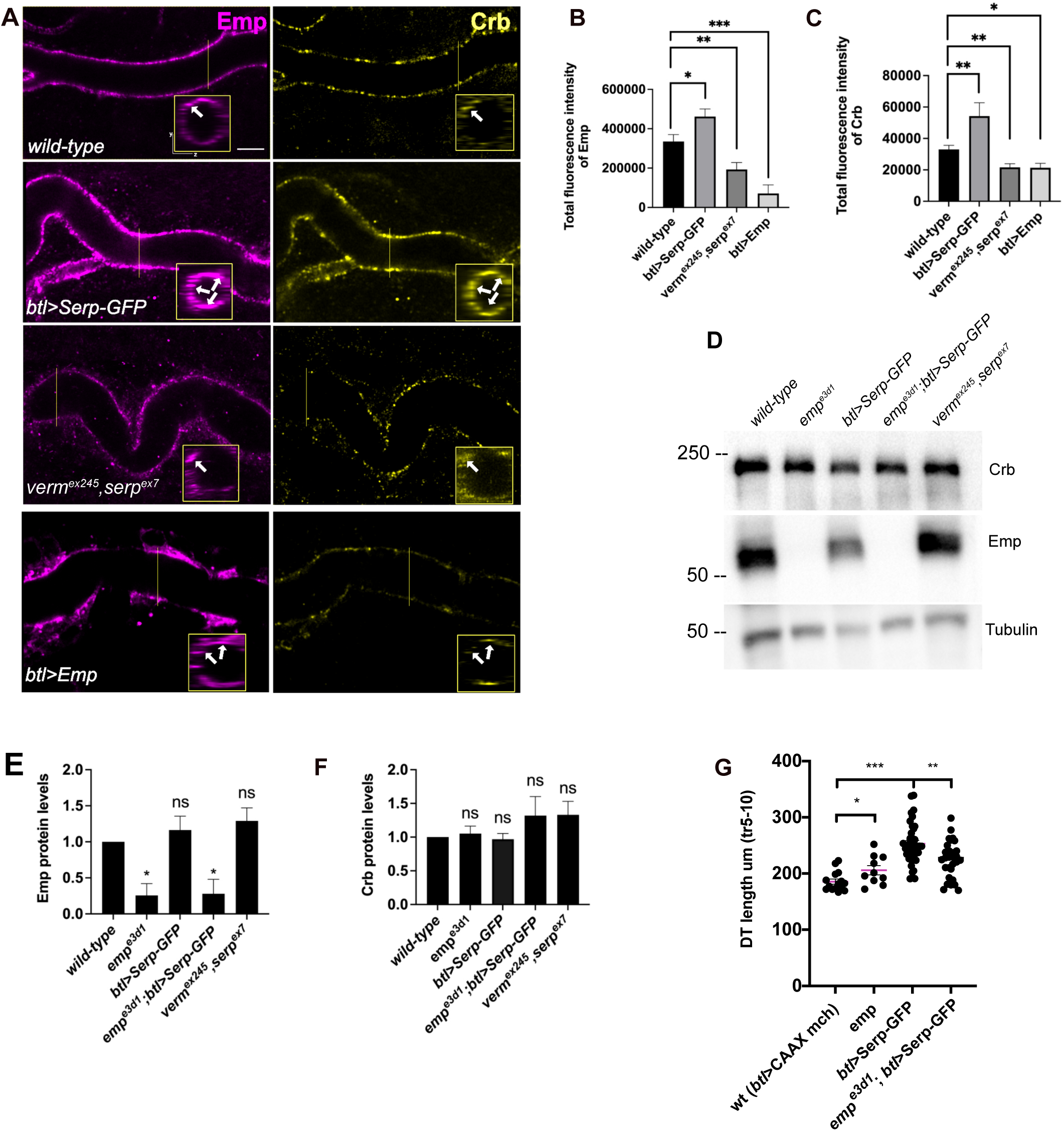
Serp overexpression induces apical Emp accumulations and tracheal over-elongation. (A) Confocal images stained for Emp and Crb, in *wild-type*, *btl>Serp-GFP*, *verm^ex245^,serp^ex7^* mutant and *btl>Emp* embryos. Inset shows zoomed view of Emp and Crb signals. The arrows indicate the accumulation of Emp and Crb in YZ plane. (B-C) Bar plots showing total fluorescence intensities of apical enriched Emp (B) or Crb (C) in *wild-type* (n = 15), *btl>Serp-GFP* (n = 12), *verm^ex245^,serp^ex7^* (n = 10) mutant and *btl>Emp* (n = 7) embryos. (D) Representative western blot from protein lysates of *wild-type*, *emp^e3d1^*, *btl>Serp-GFP*, *emp^e3d1^;btl>Serp-GFP* and *verm^ex245^,serp^ex7^* mutants, blotted for Emp and *α*-Tubulin. (E) and (F) shows the quantification of protein levels of Emp and Crb respectively, based on four independent Western blot experiments (n = 4). (G) Plots representing the length of the tracheal DT in µm (tr 5-10) from *btl>CAAX-mcherry* (control, *n* = 15), *emp ^e3d1^* (n = 10), *btl>Serp-GFP* (n = 37) and *emp ^e3d1^;btl>Serp-GFP* (n = 29) embryos. Statistical significance shown in *p*-values; p <0,0001 ****, p <0,0005 ***, p <0,01 **, p <0,05 * and p >0,05 not significant (ns) (unpaired two tailed *t* tests). Scale bars, 5μm.

### Emp controls tube morphogenesis

Crb and DE-cad trafficking underlies the anisotropic growth of the apical surface and elongation of *Drosophila* airways (Förster and Luschnig, 2012; Nelson *et al*., 2012). We first quantified the ratio of longitudinal/transverse cell junction lengths (LCJ/TCJ) in *emp* mutants and *wild-type* embryos stained for Crb. The longitudinal junction length of tracheal cells increased in *emp* mutants, while the length of junctions along the transverse tube axis was not affected (Figure 5A, B, C). The selective accumulation of the apical polarity protein Crb at longitudinal junctions relates to the extend of tube elongation. (Olivares-Castiñeira and Llimargas, 2018). Consistent with this, we detected increased Crb signals along the longitudinal but not transverse junctions in over-elongated tubes of *emp* mutants (Figure 5D-F). Stainings for the AJ component DE-cad showed increased levels along both longitudinal and transverse junctions in *emp* mutants (Figure 5D-F). To examine whether the intensity differences may be due to changes in overall protein levels, we analyzed the relative protein levels of Crb and DE-cad in *emp* mutants by western blots. We observed no significant difference in total protein levels of Crb (Figure 5G, I) or DE-cad (Figure 5H, J) in *emp* mutant embryos compared to *wild-type*. These results suggest that Crb accumulates in LCJ and DE-cad in LCJs and TCJs in *emp* mutants presumably due to defects in their endocytic uptake and trafficking. The relative LCJ vs TCJ intensities (Raw Intensity/Junctional length) of stainings for the AJs component *α*-CAT (*α*-Catenin) (Pai *et al*., 1996) and the SJs protein Dlg (disc large) (Olivares-Castiñeira and Llimargas, 2018; Sharifkhodaei, Gilbert and Auld, 2019) were indistinguishable between *emp* and *wild-type* embryos (Figure 4-figure supplement 1A-D), suggesting that intracellular junction components are intact in the mutants. Similarly, a dextran leakage assay comparing paracellular junction integrity of *wild-type*, *emp,* and *ATPα* mutant embryos showed that *emp* loss does not affect general SJ integrity (Figure 4-figure supplement 1A-D, E). Overall, these results suggest that Emp function regulates the apical membrane levels of Crb and DE-cad without majorly affecting junctional integrity or function.

**Figure 5.**
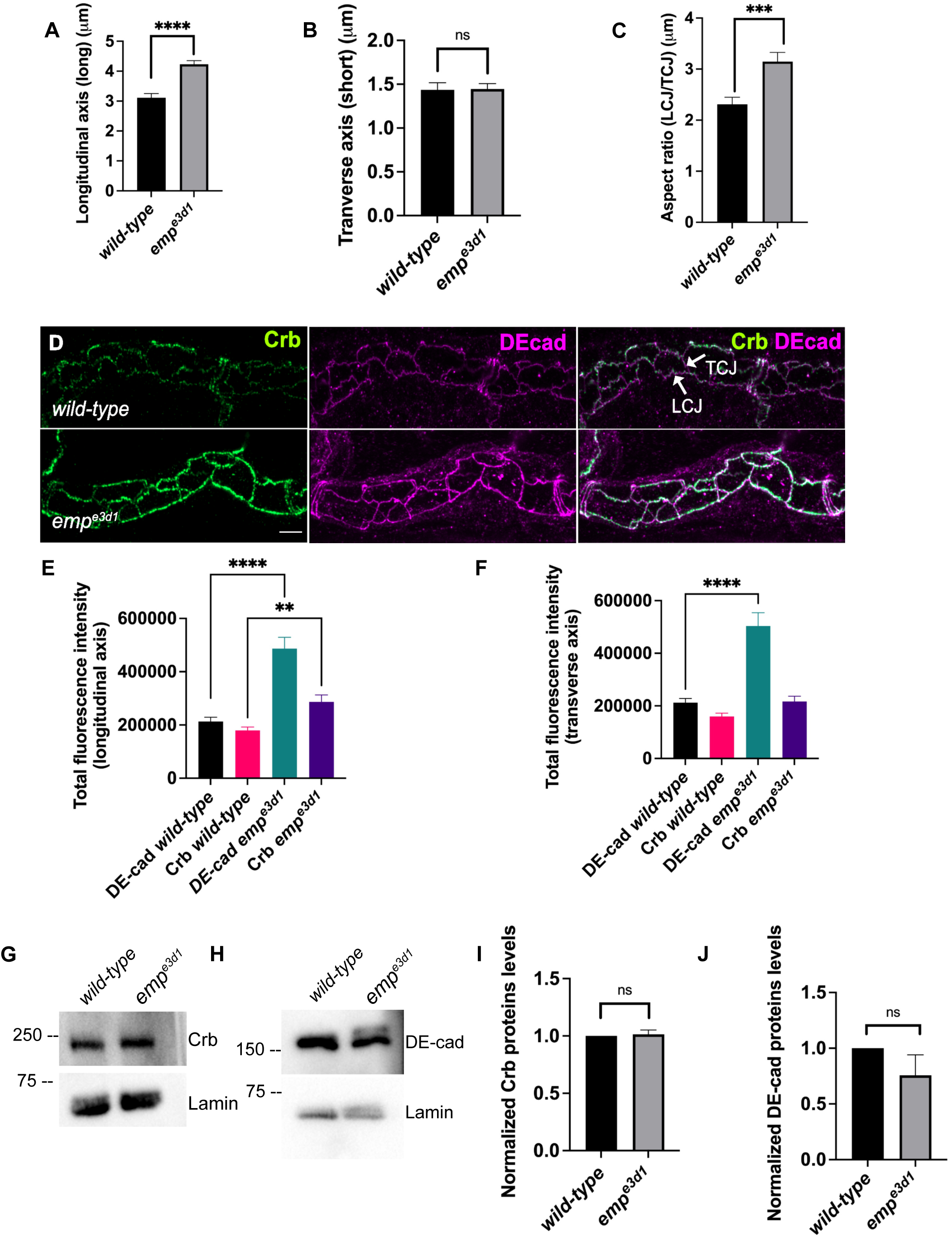
Emp modulates the Crb and DE-cad levels to control tracheal tube elongation. (A) and (B) bar plots showing the length (in *μm*) of longitudinal and transverse cell axis, respectively in *wild-type* and *emp^e3d1^* embryos. (C) Bar plots showing the aspect ratio, between longitudinal over transverse axis (LCJ/TCJ) in *wild-type* (n = 30), and *emp^e3d1^* (n = 30), mutant embryos. p < 0,0001****, p < 0,0005*** and p > 0,05 not significant (ns) (unpaired two tailed t tests). (D) Projection images of *wild-type* and *emp^e3d1^* mutant embryos, from stage 16, stained for Crb and DE-cadherin. (E) and (F) shows quantifications of the fluorescence intensities of Crb and DE-Cadherin along longitudinal (E) and transverse (F) axis in *wild-type* (n = 78), and *emp^e3d1^* (n = 66), mutants. p < 0,0001**** and p < 0,01** (Mann-Whitney test). (G - H) Representative western blot from protein lysates of *wild-type* and *emp^e3d1^* mutants, blotted with anti-Crb (G) or anti-DE-cad (H) and anti-Lamin (control). (I) and (J) are quantifications of Crb and DE-cad protein levels, respectively, based on three independent western blot experiments (n = 3). Statistical significance shown in p-values; p > 0,05 not significant (ns) (unpaired two tailed t tests). Scale bars, 5μm. **Figure 5—source data 1** This zip archive contains the raw unedited western-blot shown in Figure 5G, H.

### Emp modulates the apical actin organization

Our analysis until now suggests that Emp is a scavenger receptor involved in the continuous, basal-level trafficking of Serp during tube elongation and in its massive uptake during luminal protein clearance. Since the organization of the transverse actin bundles restrict both tube elongation and the timing of luminal protein clearance, we investigated the apical actin cytoskeleton in *emp* mutants. We first examined the apical F-actin in DTs of late-stage embryos by live imaging using *btl>Moe-GFP*. At 17,5h AEL, at the time interval of airway protein clearance, *emp* mutants showed a more diffuse and continuous apical actin bundle density compared to *wild-type* embryos (Figure 6A, B). We also detected higher apical accumulation of the dDAAM formin, in *emp* mutants compared with *wild-type* embryos by antibody stainings. These observations suggested that Emp modulates the actin bundle organization, presumably upon engagement with its luminal cargoes.

**Figure 6.**
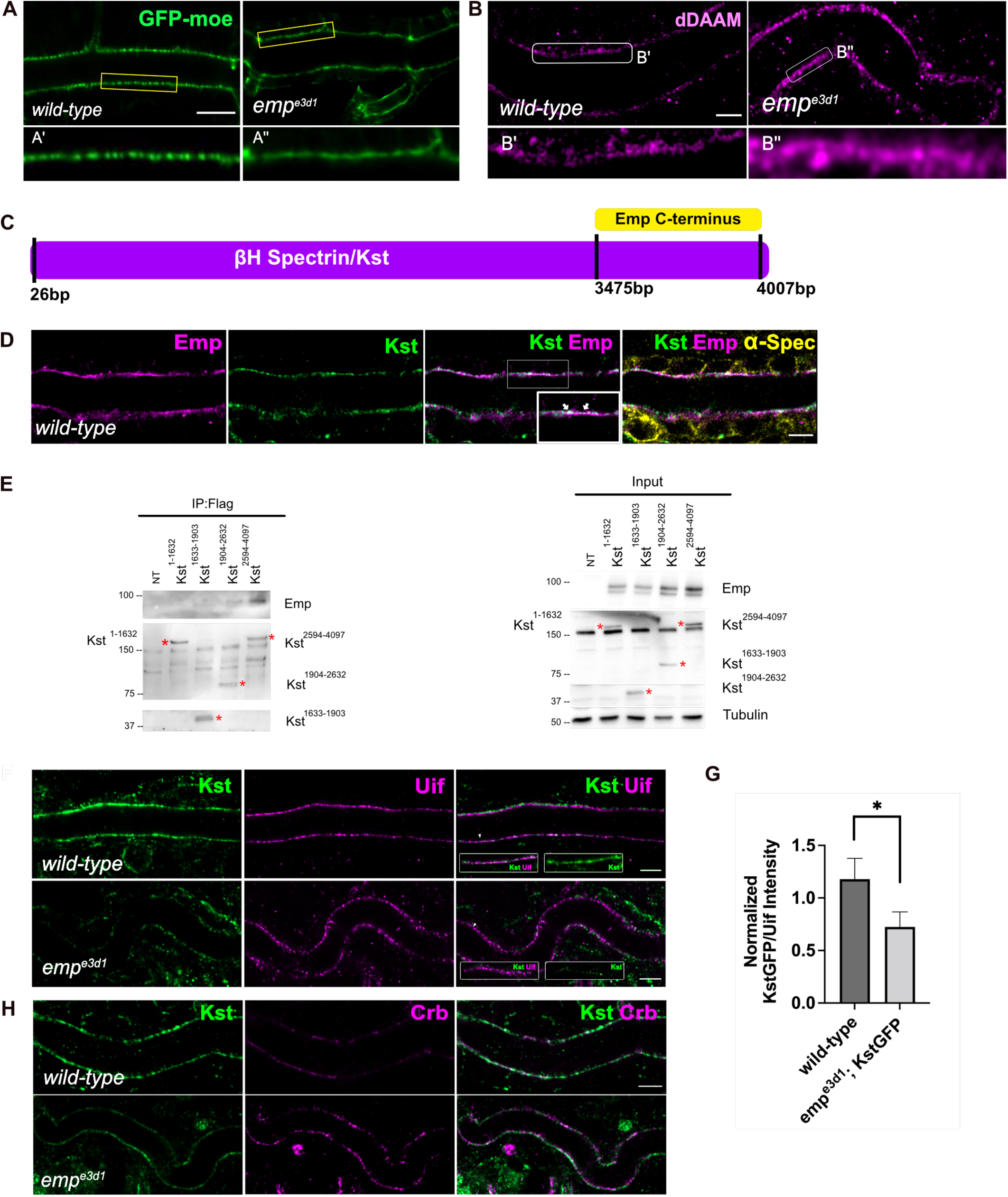
Loss of Emp affects apical F-actin organization. (A) Confocal images showing live DT of *wild-type and emp^e3d1^* mutant embryos from 17,5h AEL expressing the actin reporter *btl>moe-GFP*. (A’, A’’) zoomed views of areas indicated by the rectangular frames of (A) image. (B) Confocal images of dDAAM stainings in *wild-type* and *emp^e3d1^* mutant embryos. (B’, B’’) shows magnified regions of (B), indicated with white rectangle. (C) Schematic view of the interaction domains of Emp C-terminus (484aa to 520aa) and *β*HSpectrin/Kst, as obtained by the Y2H screen. (D) Confocal images showing the DT of Kst-Venus expressing embryos (*wild-type*) stained for Emp, GFP (kst), and *α*-Spec (*α*-Spectrin). (E) Co-immunoprecipitation of Flag-tagged Kst constructs from transfected S2 cells lysates, blotted with anti-Emp and anti-Flag. Input 2% is indicated. (F) Confocal images showing endogenous Kst-Venus in the DT of *wild-type* and *emp^e3d1^* mutants, stained for GFP (kst) and Uif (Uninflatable). (G) Bar plot showing Kst-GFP levels in *wild-type* (n = 5), and *emp^e3d1^* (n = 5), mutant embryos in relation to Uif. (H) Confocal images showing *wild-type* and *emp^e3d1^* mutant embryos stained for GFP (kst) and Crb. Statistical significance shown in p-values, p <0,0005***, p <0,05* and p >0,05 (ns) (unpaired two tailed *t* tests). Scale bars, 5μm, (B, D, F, H) and 10μm (A).

To identify proteins that might provide a direct connection between the cytoskeleton and Emp, we used the cytoplasmic C-terminus of Emp as bait in a yeast two hybrid (Y2H) screen at Hybrigenics. This uncovered 3 potential interacting proteins, CG32506 encoding a RabGAP protein, CG34376 encoding a protein with a predicted Zinc Finger motif and βH-Spec/Kst (Figure 6C). We further characterized the interaction with βH-Spec/Kst because of its established role in the apical cytoskeleton organization in *Drosophila* epithelial tissues (Thomas and Williams, 1999; Phillips and Thomas, 2006). The *α*- and *β*-heavy Spectrin subunits form tetramers, which assemble in two dimensional networks together with actin underneath the apical plasma membranes. Additionally, βH-Spec/Kst is required for the early steps and endocytic trafficking of V-ATPase in the brush border of intestinal epithelial cells (Phillips and Thomas, 2006). We used a knock-in, fusion construct of Venus into the *kst* locus to detect endogenous, βH-Spec/Kst together with Emp and α-spectrin in epithelial cells of the trachea and hindgut. As expected Kst-Venus colocalized with Emp apically, whereas α-spectrin was also detected along the lateral sides. To confirm the Emp binding to Kst and also map their interaction domains we used immunoprecipitation experiments of tagged proteins in *Drosophila* S2 cells. We expressed V5-tagged Emp and a series of constructs expressing different fragments of βH-Spec/Kst protein fused to the FLAG epitope. We found that the intracellular C-terminus of Emp co-precipitates with the C-terminal region of βH-Spec/Kst (Figure 6E) consistent with their interaction detected in the yeast-two-hybrid system. Additionally *kst^2^* mutants show similar tracheal over elongation phenotypes with *emp* suggesting a functional interaction between Emp and Kst (Figure 6-figure supplement 1A, B). To further test this, we analyzed the localization and abundance of tagged βH-Spec/Kst in the airways of *emp* mutants. The apical levels of βH-Spec/Kst were severely reduced, while Crb staining was increased and staining of an unrelated apical protein Uninflatable remained unaffected in the *emp* mutants compared to the *wild-type* (Figure 6F-H). The localization or levels of Emp were not noticeably affected in *kst* mutants (Figure 6-figure supplement 1C). Together, these results indicate that the C-terminal intracellular domain of Emp binds to apical βH-Spectrin (Kst) and controls the spectrin cytoskeleton presumably by stabilizing βH-Spec/Kst. Additionally, in *emp* mutants the intensity and distribution of actin bundles is distorted and the levels of the diaphanous-related formin, DAAM are increased.

### Emp regulates Src phosphorylation

Src phosphorylation and activation is required for tracheal tube elongation and controls Crb accumulation at the longitudinal cell junctions (Olivares-Castiñeira and Llimargas, 2018), and the Beitel laboratory proposed that Src together with DAAM orient the directed membrane expansion during tube elongation (Nelson *et al*., 2012). Since *emp* mutants showed higher Crb accumulation in longitudinal junctions and an overall increase in the apical levels of DAAM, we compared Src phosphorylation (p-Src) levels by immunostainings and by western blot in *wild-type*, *emp* mutants, *btl>Serp-GFP* and *emp;btl>Serp-GFP* embryos (Figure 7A-C). The specificity of the p-Src^419^ antibody was first confirmed by staining of *src42A src64B* double mutants (Figure 7-figure supplement 1A). Both *in situ* staining and the western blot analysis showed increased p-Src levels in *emp* mutants compared to *wild-type* (Figure 7A-C). Interestingly, p-Src levels were also increased in the *verm serp* double mutant embryos and decreased in *wild-type* embryos overexpressing *btl>*Serp*-GFP* (Figure 7B). This decrease by Serp overexpression was partly ameliorated in *emp* mutants (Figure 7A-C), suggesting that the effect of Serp-GFP overexpression on Src phosphorylation is, at least partly, mediated by Emp on the cell surface. We also detected increased p-Src levels in total protein extract from *emp* and *verm serp* mutant embryos (Figure 7B) suggesting that their interaction controls Src phosphorylation in other ectodermal tissues. In agreement with this, neurons of the ventral neve cord in *emp* mutant embryos also sowed increased p-Src levels compared to *wild-type* (Figure 7-figure supplement 1B). To test if the increase in p-Src levels in *emp* mutants underlies the apical membrane over-elongation defects, we performed genetic interaction experiments between *src42A* and *emp* mutants. As expected, *emp* mutants showed over-elongated tubes while the *src42A^E1^* mutants showed short tubes (Nelson *et al*., 2012). The increase of tube length in *emp* mutants was significantly suppressed in *emp;src42A^E1^* embryos, where the levels of Src protein were reduced (Figure 7D, E). Similarly, the increased apical accumulation of Crb and DE-cad in *emp* mutants was partly restored by overexpression of a *Src42A^DN^* dominant negative form (*Src42A^DN^*) in emp mutant embryos. (Figure 7-figure supplement 1C-F). These results suggest that downregulation of src activity and the levels of luminal Serp-GFP are sensed by Emp to control apical membrane protein endocytosis and trafficking, and thereby elongation.

**Figure 7.**
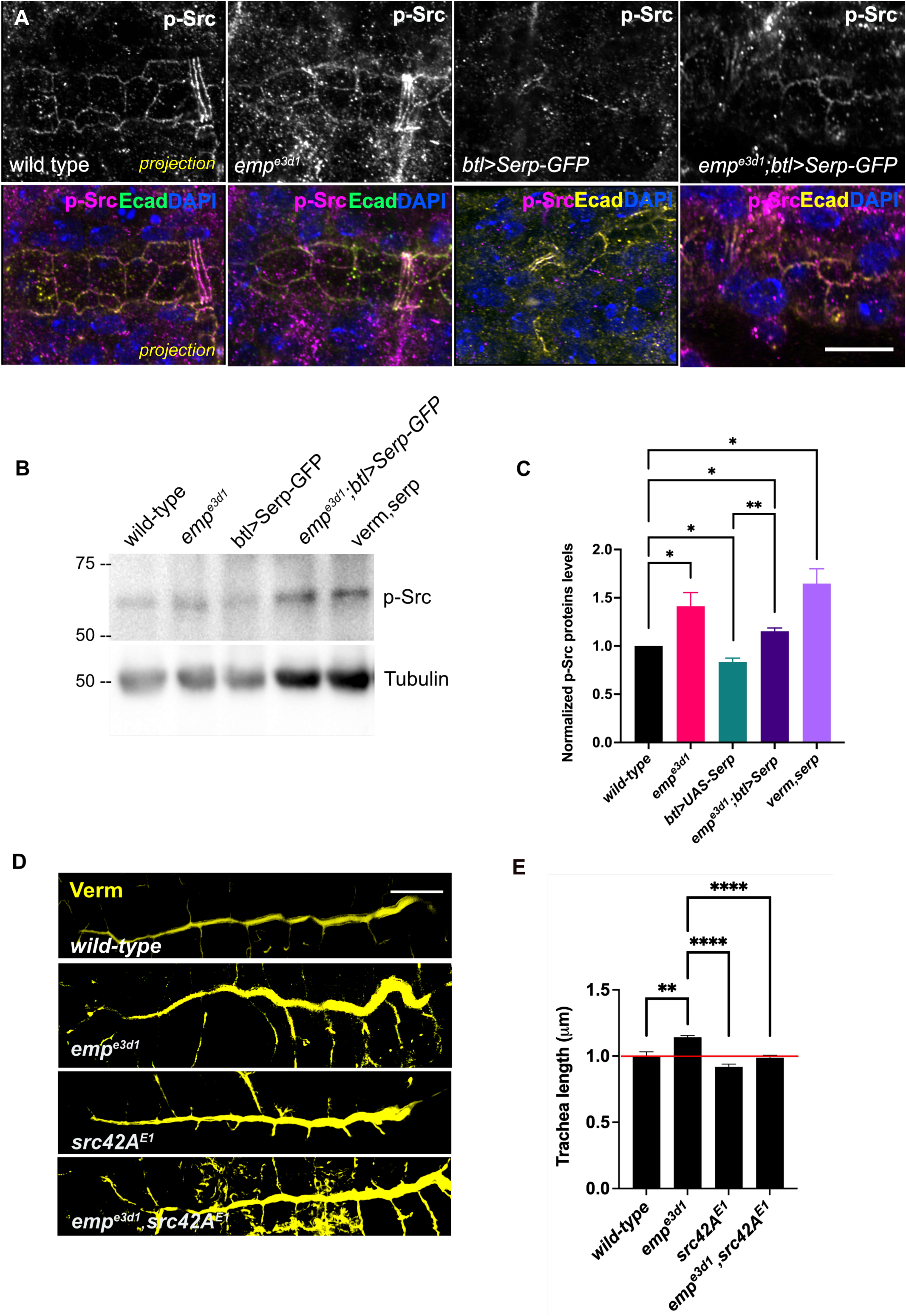
Elevated p-Src levels in *emp^e3d1^* mutants. (A) Confocal images showing projection of the tracheal DT of *wild-type*, *emp^e3d1^* mutants, *btl>Serp-GFP* and *emp^e3d1^;btl>Serp-GFP embryos* at late stage 16 to early stage 17, stained for endogenous p-Src. (B) Representative western blot from protein lysates of *wild-type*, *emp^e3d1^* mutants, *btl>Serp-GFP*, *emp^e3d1^;btl>Serp-GFP* and *verm,serp* double mutant embryos, blotted with anti-p-Src and anti-⍺-Tubulin. (C) Quantifications of p-Src protein levels based on three independent Western-blot experiments (n = 3). (D) Representative images of tracheal length size (DT), in wild-type, empe3d1, src42AE1 and empe3d1,src42AE1 embryos stained for the luminal marker Verm. (E) Plots show the quantification of tracheal DT length of *wild-type* (n = 6), *emp^e3d1^* (n = 7), *src42A^E1^* (n=10) and *emp^e3d1^;src42A^E1^* (n=7) embryos. Error bars denotes s.e.m. wild-type, p < 0,005**, and p < 0,0001****(unpaired two tailed *t* tests). Scale bars, 10 μm and 50μm for images (A) and (D) respectively.

## Discussion

Emp, a CD36 homologue, is a selective scavenger receptor required for endocytosis of a subset of luminal proteins. The endocytosis defects in *emp* mutants become most apparent during the massive endocytosis wave that removes all secreted luminal components just before gas filling of the airways. Our comparison of the endocytosis requirements of Serp and Gasp together with the LDLr-domain swap experiments defined a subset of Emp cargoes and indicates that the LDLr-domain targets cargoes to Emp through clathrin-independent endocytosis. These selective requirements of Serp and Gasp internalization are consistent with the view that the choice of endocytic route is a cargo-driven process (Mettlen *et al*., 2018).

The machinery involved in class B scavenger receptor endocytosis have not been characterized in *Drosophila*, but an important characteristic of all clathrin-independent endocytosis pathways is their dependance on the dynamic control of actin polymerization to distort the plasma membrane for cargo internalization (Mayor, Parton, and Donaldson 2014). Pioneering biochemical experiments suggested that CD36 receptor clustering is essential for its internalization and signaling in response to multivalent cargo binding. Single molecule tracking of CD36 in primary human macrophages showed that the un-ligated receptor diffuses in linear confinement tracks set by the actin cytoskeleton. These diffusion tracks enable clustering and internalization upon oxLDL-ligand addition (Jaqaman *et al*., 2011). Similarly, in human endothelial cells the actin cytoskeleton is required for the increase of CD36 clustering upon thrombospondin binding and Fyn, a Src-family kinase activation (Githaka *et al*., 2016). These studies argue that CD36 is confined in cytoskeletal tracks and cargo/ligand binding induces its clustering and signaling ability. Similarly to CD36, Emp function is also tightly connected with the apical cytoskeleton. First, *emp* activity is confined in apical “macro”-domains along the longitudinal tube axis by the transverse, DAAM-dependent actin filaments. Disruption of these filaments induces massive endocytosis and re-localization of Emp. Additionally, the initially punctate Emp distribution along the apical membrane becomes aggregated upon overexpression of luminal Serp, suggesting that LDLr-domains on chitin-binding proteins induce Emp clustering. Apart from the Emp similarities to CD36, our results also reveal an unexpected direct function of Emp in organizing the apical Spectrin and actin cytoskeleton. The distribution of the DAAM-formin, the transverse actin bundle density and the apical accumulation of Kst are all disrupted in *emp* mutants. Because we showed that the conserved C-terminal intracellular domain of Emp binds directly to Kst, we propose that a central function of Emp, and possibly CD36, is to also directly organize the epithelial cytoskeleton. This notion is supported by the ability of human CD36 to partially rescue the *emp* mutant phenotypes when overexpressed in the *Drosophila* airways.

How can a scavenger receptor restrict epithelial tube elongation? In the interval of embryonic stages 13 to 16, regulated Src and DAAM activities promote tube elongation and the formation and alignment of the transverse actin bundles. These bundles inhibit Emp internalization and restrict Serp endocytosis and trafficking to the longitudinal tube axis. This suggests that endocytosis of luminal proteins during tube elongation is controlled by at least 2 parallel pathways, one enabling Emp-endocytosis along the longitudinal tube axis and one restricting it along the transverse axis. In the bundle-free membrane domains, Emp associates with Kst and establishes an endocytosis domain, where Serp mediated Emp clustering leads to internalization (Figure 8 A, B). A similar function for Kst enabling endocytosis and trafficking of apical H^+^ V-ATPase has been proposed in the brush border of the larval *Drosophila* intestine, where its anchoring to the membrane remains unknown. The luminal levels of chitin deacetylases, like Serp, and other chitin modifying proteins are transcriptionally regulated (Yao *et al*., 2017) and are predicted to control the biophysical properties of the apical ECM (Cui, Yu and Lau, 2016). We infer that luminal Serp bound to chitin generates a multivalent ligand and is continuously recognized and endocytosed by Emp selectively along the longitudinal tube axis. Together with the clustered receptors, apical membrane and “passenger” transmembrane proteins are expected to follow in the Emp endocytic vesicles. The over-elongation defects in *emp* mutant airways can be explained by the failure to balance elongation induced by Src activation with endocytosis of membrane and transmembrane regulators on the longitudinal tube axis (Figure 8 A, B).

**Figure 8.**
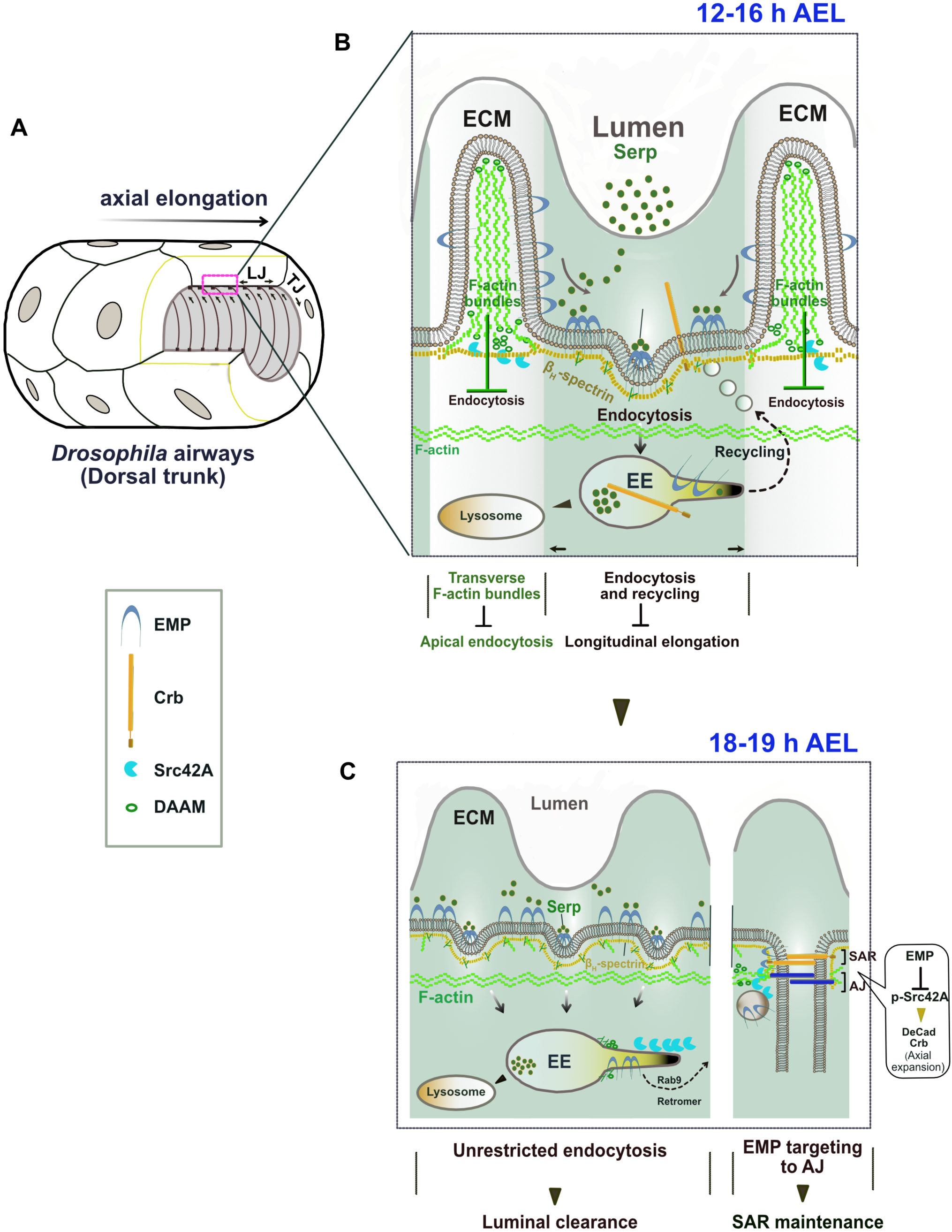
The proposed model of Emp regulation. (A) 3D drawing shows the multicellular tube (Dorsal Trunk) of *Drosophila* airways. The annular ridges of the apical ECM are indicated in relation to the longitudinal elongation. LJ: Longitudinal junctions; TJ: transverse junctions. (B) Schematic zoom of the apical ECM region (A) showing the formation of two apical domains in the membrane-cytoskeleton interface of tracheal cells at 12-16h AEL. Transverse F-actin bundles (white color zone) restrict apical Emp endocytosis along the transverse tube axis. In F-actin bundle-free membrane region (green color zone), Emp is associated with *β*H-Spectrin and Serp promotes Emp clustering leading to endocytosis and recycling of Serp and others “passenger” transmembrane proteins (i.e. Crb) along the longitudinal tube axis. (C) Drawing of the apical, SAR and AJ regions during luminal protein clearance (18-19h). Emp clears luminal Serp by endocytosis and trans-locates to the SAR/AJ to restrict p-Src42 activity and to control DE-Cad, Crb levels at the SAR/AJ.

At the initiation of protein clearance, the transverse bundles are transiently resolved and luminal proteins together with apical transmembrane proteins and membrane are massively internalized and targeted for degradation or recycling to the junctional areas (Figure 8C). The mechanism of Crb recycling in airway cells involves retromer components (Olivares-Castiñeira and Llimargas, 2017) but the selective targeting mechanism to longitudinal junctions is still unknown. The re-localization of Emp to the apical epithelial junctions during airway maturation is accompanied by massive alteration in Src activity, which is not restricted to the tracheal system but is common to several ectodermal organs including neurons. This suggests that scavenger receptor endocytosis has a general, but poorly understood role, in embryonic morphogenesis of epithelial tissues. Our work on Emp provides an entry towards elucidating the roles of scavenger receptor class B in development and pathogenesis.

## Material and Methods

### Drosophila strains

The *emp^e3d1^* null mutant was generated by FLP-FRT site-directed recombination using two piggyBac elements (PBacWH#021071 and PBacRB#e0441541). Embryos trans-heterozygous for Df(2R)BSC08 and *emp* were lethal showing identical phenotypes to *emp* homozygous embryos. The list of strains used are list in Supplementary Table 1. w^1118^ was used as the *wild-type* strain. In all experiments CyO and TM3 balancer strains carrying dfd-GFP were used to identify the desired genotypes. Flies were raised at 25 °C and 50% humidity, with a 12 hrs light–dark cycle.

### Molecular Biology and Transgenic flies

Complementary DNA (cDNA) encoding for *Human CD36 (*RC221976) was cloned using Hifi DNA assembly kit (NEB E5520S) into the pJFRC-MUH vector (Addgene, plasmid 26213). bglII and XbaI were used to clone the amplified PCR fragment. *UAS-Emp* transgene was generated by cloning the cDNA sequencing of CG2727 by PCR amplification with primers containing EcoRI and KpnI. The *UAS-Gasp* construct was generated by sequential cloning of Gasp cDNA using primers containing EcoRI and XhoI. Further, LDLr domain and mCherry was subcloned using enzymes XhoI and XbaI. The *UAS-Serp^FL^-GFP* and *UAS-Serp^LDLr^-GFP* constructs was generated according to *Shenqiu Wang et al. Current Biology 2006*, and the *UAS-Serp^CBD^-GFP* was provide S. Luschning. *mCherry-Emp-V5His* construct was cloned using Hifi DNA assembly kit (NEB E5520S into the pAc5.1/V5-His A vector (Invitrogen) using the following enzymes Acc65I and XhoI. The diferent Kst constructs were generously provided by N. Tapon. All the plasmids were confirmed by sequencing. Polyclonal antibody (anti-Emp) obtained by immunization with bacterially expressed recombinant polypeptides corresponding to amino acids 46-460aa of Emp-A. Anti-sera obtained from immunized rats (Genscript).

### Co-Immunoprecipitation and western blot analysis

*Drosophila* S2 cell extracts and Co-IP were prepared as previously described (Tsarouhas *et al*., 2019) and (Fletcher *et al*., 2015) respectively. The FLAG-tagged Kst constructs were provided by Nic Tapon (Fletcher *et al*., 2015). For detection of purified proteins and associated complexes, ChemiDoc XRS + system (BioRad) was used. Western blots were probed with mouse anti-FLAG M2 (1:3000, Sigma, F3165), rat anti-Emp and rabbit anti-α-tubulin (1:2000, Cell Signaling, 11H10). For western blot analysis, *Drosophila* embryos were collected 12–20 h AEL and lysed in 20 μl of lysis buffer containing 50 mM HEPES (PH 7.6), 1 mM MgCl_2_, 1 mM EGTA, 50 mM KCl, 1% NP40, Protease inhibitor cocktail tablets (Roche #11697498001) and Phosphatase inhibitor cocktail 2 (Sigma Aldrich #P5726). The lysates were centrifuged at maximum speed (30,060 × *g*) for 10 min at 4 °C. Protein loading buffer (50 mM Tris/HCl, pH 6.8, 2% sodium dodecyl sulfate (SDS), 5% glycerol, 0.002% bromophenol blue) was added to the supernatant and samples were analyzed by SDS-polyacrylamide gel electrophoresis (PAGE) and immunoblotting according to standard protocols, using the ChemiDoc XRS + system (BioRad), after application of the SuperSignal West Femto Maximum Sensitivity Substrate (ThermoFisher Scientific, 34096). The following primary antibodies were used at the indicated dilutions: rabbit anti-α-tubulin (1:2000, Cell Signaling, 11H10), rabbit anti-Phospho-Src (1:750 Tyr418, ThermoFisher) and Lamin ADL195 (1:100, DSHB).

### Quantification of western blots

For the western blot analysis, the actual signal intensity of each band of interest was estimated after subtraction of the background using ImageJ/Fiji software. The values were then divided by the corresponding intensities of the loading control (α-tubulin or lamin).

### Immunostaining

Embryos were dechorionated in 5% bleach and fixed for 20 min in 4% formaldehyde saturated in heptane as described in (Patel, 1994). The following antibodies were used: mouse anti-Ptp10D (1:10, 8B22F5, Developmental Studies Hybridoma Bank, DSHB), rabbit anti Phospho-Src (1:400, Tyr 419, Thermo-Fisher), mouse anti Dlg (1:100, 4F3 DSHB), mouse anti-Crb (1:10, Cq4 DSHB), mouse anti-Coracle (1:100, C615.16 DSHB), DAAM antibody was a kind gift from József Mihály, rabbit anti-GFP (1:400, A11122, Thermo-Scientific), chicken anti-GFP (1:400, abl3970, Abcam), mouse GFP (1:200, JL-8, Clontech) gp anti-Verm (Wang *et al*., 2006), gp anti-Gasp (Tiklová, Tsarouhas and Samakovlis, 2013), mouse anti-Flag M2 (1:3000, Sigma, F3165), Serp antibody were provided by S.Luschning. Secondary antibodies conjugated to Cy3 or Cy5 or Alexa Fluor-488 and -568 (Jackson Immunochemicals) were used and diluted as recommended by the manufacturer. For rat anti-DE-Cad (1:50, DSHB) embryos were fixed with 4% PFA – heptane for 20min. Embryos expressing *moeGFP* were dechorionated, devitellinized by hand and fixed in 4% formaldehyde (methanol free) in PBS–heptane for 20 min. Stained embryos were imaged with an Airy-scan-equipped confocal microscope system (Zeiss LSM 800, Carl Zeiss) using a Plan-Apo 63x/1.40 DIC oil immersion objective.

### Yeast two-hybrid screen

The screen was carried out by Hybrigenics using a prey library constructed from RNA of embryos that were 0–24-h old. A fragment encoding the C-terminus domain of Emp (amino acids 484–520) was inserted into the pB27 vector (N-LexA–bait-C fusion) and was used to screen 167 million clones.

### qPCR

Embryos dechorionated in bleach, hand-sorted for GFP expression, collected in 300 uL TRizol LS Reagent and stored at −80°C until further use. For RNA extraction, the embryos were homogenized in TRIzol LS Reagent using a 1.5 mL tube pestle and the total RNA was purified using the Direct-zol RNA MicroPrep kit (R2060, Zymo Research). RNA was resuspended in RNase-free water and subsequently treated with DNAse I (AMPD1-1KT, Merk), for genomic DNA removal. Then 400 ng of RNA was reverse transcribed using High-Capacity RNA to cDNA kit (4387406 ThermoFisher). The cDNA products were subsequently diluted 1:5 and 2 μl were used as a template in each qPCR reaction. qPCR was performed using iTaq Universal SYBR Green Supermix (Biorad). Generation of specific PCR products was confirmed by melting-curve analysis. Ct values for all genes were normalized to the levels of *Rp49*. For data analysis, the delta-delta Ct values was applied. The sequences of primers used are provided in Supplementary Table 2.

### Live imaging

Dechorionated embryos mounted in a glass-slide with a gas permeable membrane (Tsarouhas *et al*., 2019). Widefield live imaging performed to analyze protein clearance and gas filling on embryos as described in (Tsarouhas *et al*., 2019). For confocal live-imaging, embryos were imaged with a scanning confocal microscope (LSM 780, or 800 Carl Zeiss) equipped with an Argon and an HeNe 633 laser using a C-Apochromat ×63/1.2 NA water objective. *Z*-stacks with a step size of 0.5–1.0 μm were taken every 6 min over a 3–8-h period. For high-resolution confocal live imaging, an Airy-scan-equipped confocal microscope system (Zeiss LSM 800, Carl Zeiss) was used. *Z*-stacks (0.16–0.2-μm step size) were taken every 15 min over a 2–4-h period using a Plan-Apo 63x/1.40 DIC oil or a C-Apochromat ×63/1.2 NA water objective (Zeiss). Raw data were processed with the Airy-scan processing tool available on the Zen Black software version 2.3 (Carl Zeiss). Images were converted to tiff format using the Zen Black or ImageJ/Fiji software.

### Morphometric analysis

Tube length measurements were conducted in embryos stained for the luminal markers Verm or Serp. Tracheal lengths were measured by tracing the length of the dorsal trunk determined by the luminal markers using the freehand line selection tool of ImageJ/Fiji software. For metamere length measurements, we traced the DT length between the corresponding TC (transverse connective) branches. Longitudinal and transverse cell junctions were defined according to the angles from the DT axis (angle = 0°). Junctions with clear orientation angle 0°±30° or 90° ±30° were defined as longitudinal or transverse, respectively. The rate of gas-filling calculated as the percentage of embryos with gas-filled tracheae divided by the total number of embryos analyzed.

### Dextran injections

For dye-permeability assays, 10 kDa Dextran-TR (ThermoFisher Scientific) was injected into late-stage 16 embryos (after the maturation of SJ) as described in ref (Jayaram *et al*., 2008). Injections were performed in a microinjection system (FemtoJet, Eppendorf) coupled to an inverted fluorescence microscope (Cell Observer, Zeiss).

### Quantification of fluorescence intensity

The total fluorescence intensity signal of Emp, Crb, DE-cad, Dlg and *α*-Cat was measured with ImageJ by manually drawing a 5-pixel line with the “Freehand Line tool”, over the junctional region of the tracheal cells. The total fluorescence intensity in TCJ and LCJ of Crb and DE-cad was measured according to (Olivares-Castiñeira and Llimargas, 2018). Background signals of 5-pixel line, were subtracted from the intensities. For the quantification of fluorescent intensity on apical Emp positive puncta, 0,75 μm^2^ squares on each puncture were defined as the regions of interests (ROIs). Mean fluorescent intensity within these ROIs was measured in Fiji. These values were divided by the corresponding fluorescence intensity observed in the ROI of the co-stained Crb puncta. Background signals of 0,75 μm^2^ square-ROIs were subtracted from the intensities. Data collected from ImageJ were transferred to an Excel file for further analysis and the plotting using Graph Pad Prism.

### Statistical analysis

Statistical analysis was carried out using two-tailed *t* test for unpaired variables unless indicated. The type of statistical test, *n* values and *P* values are all provided in the figure legends. The experiments were replicated 3 - 6 times. All statistical analyses were performed using Graph Pad Prism 9.1. The number of biological replicates for all the experiments is indicated in the figure legends. No explicit power analysis was used to estimate sample sizes for each experiment.

## Acknowledgements

We would like to thank Stefan Luschnig, Stefano De Renzis, Nic Tapon, the Bloomington Drosophila Stock Center, the Drosophila Genomics Resource Center (DGRC; IN) and the Developmental Studies Hybridoma Bank (DSHB; IA) for fly strains, clones and antibodies. We thank the fly community that isolated, characterized or distributed mutant strains or antibodies. Special thanks to Flybase for the Drosophila genomic resources. We thank the Stockholm University Imaging Facility (IFSU). We thank the former ERASMUS student Claudia’s Ctortecka, members of the M. Mannervik, C. Samakovlis (particularly Ryo Matsuda), Q. Dai, S. Åström and Y. Engström laboratories for comments and support during this project. This work was funded by the Swedish Research Council and the Swedish Cancer Society to C.S; by the Magn. Bergvalls stiftelse and O. E. och Edla Johanssons vetenskapliga stiftelse to V.T; and by the German Research Foundation (DFG), grant KFO309 (project number 284237345) to C.S.

## Author contributions

A.P. designed and executed most of the experiments, analyzed most of the data and wrote the paper. V.T. designed and executed experiments, analyzed data and wrote the paper. K.S. conceived and initiated the project by generating the *emp* mutants. B.A. generated Emp-GFP construct. C.S. conceived the project, proposed experiments, analyzed data, and wrote the paper.

## Data availability

All data generated or analyzed during this study are included in the manuscript and supporting files. Unprocessed Western blots are provided as source data files in zip format.

---

**Figure 1—figure supplement 1.**
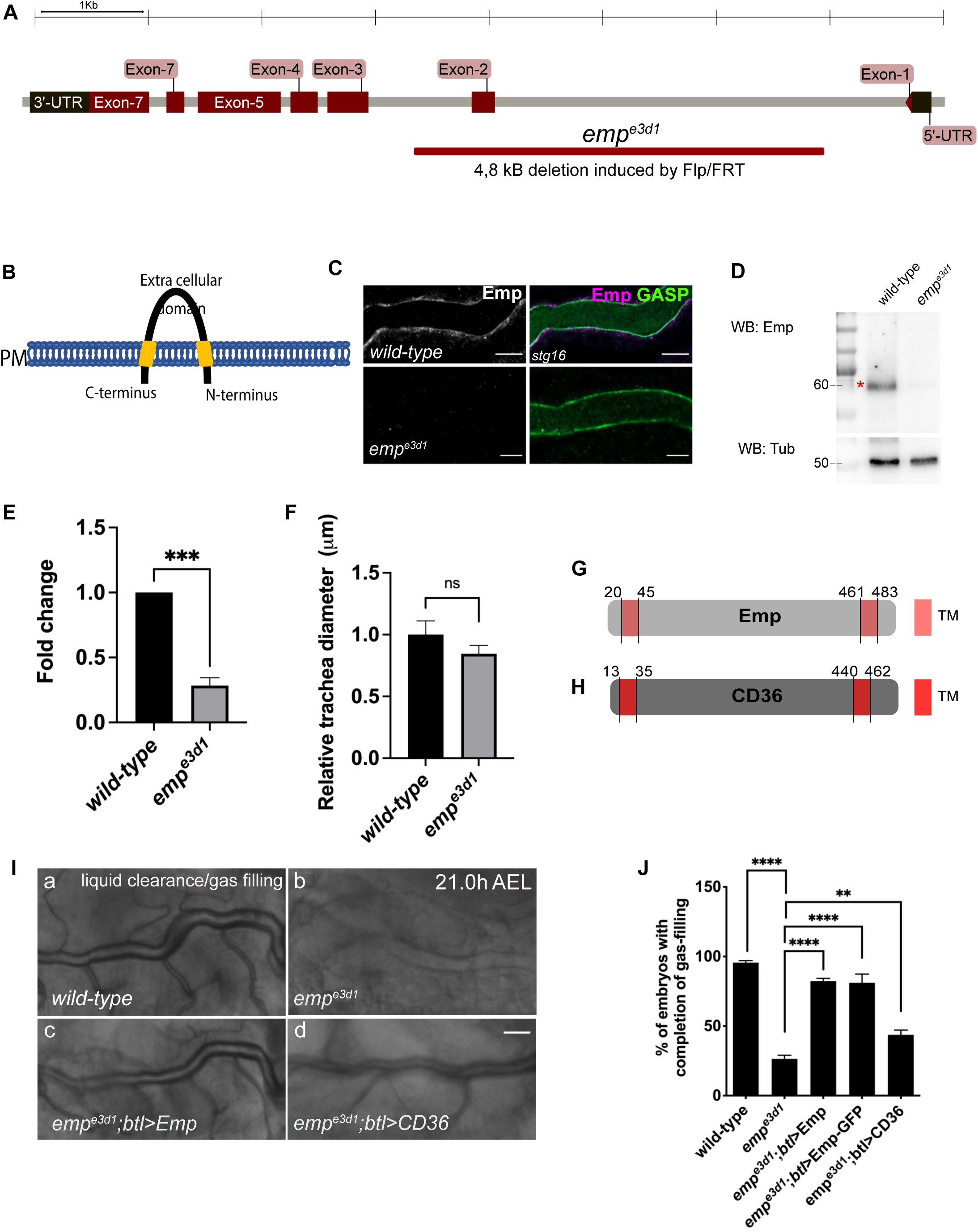
Conserved function of Emp and it’s human homologue CD36. (A) Schematic view of the *emp* (CG2727) locus. The Flp/FRT-induced 4,8Kb deletion in *emp^e3d1^*. (B) The extra cellular domain of the Emp protein selected to generate the anti-Emp antibody. (C) Confocal images of DT, showing the Emp antibody specificity. (D) Western Blot from *wild-type* and *emp^e3d1^* lysates embryos, probed for Emp and ⍺-Tubulin. (E) The expression levels of *emp* mRNA were calculated by qRT-PCR from *wild-type* or *emp^e3d1^* embryos at stage 16. (F) Bar plots showing the relative tracheal diameter of *wild-type* (n=10) and *emp^e3d1^* (n=8) embryos. (G-H) Graphical illustration of the Emp (G) and human CD36 (H) protein domains. (I) Widefield images of DT at late stage 17 of living *wild-type, emp^e3d1^, emp^e3d1^; btl>Emp* and *emp^e3d1^; btl>CD36* embryos showing gas-filing phenotype. (J) Bar graph displaying the percentage of embryos that fill with gas in *wild-type* (n=177), *emp^e3d1^* (n=182), *emp^e3d1^; btl>Emp* (n=119), *emp^e3d1^;btl>Emp-GFP* (n=178), and *emp^e3d1^; btl>CD36* (n=70) embryos. Error bars denotes s.e.m., p >0,05 not significant (ns), p < 0,005**, p < 0,0005*** and p < 0,0001**** (unpaired two tailed t tests). Scale bars, 5 μm (C) 50 μm (I). **Figure 1—figure supplement 1- source data 1** This zip archive contains the raw unedited western-blot shown in Figure 1—figure supplement 1.

**Figure 2—figure supplement 1.**
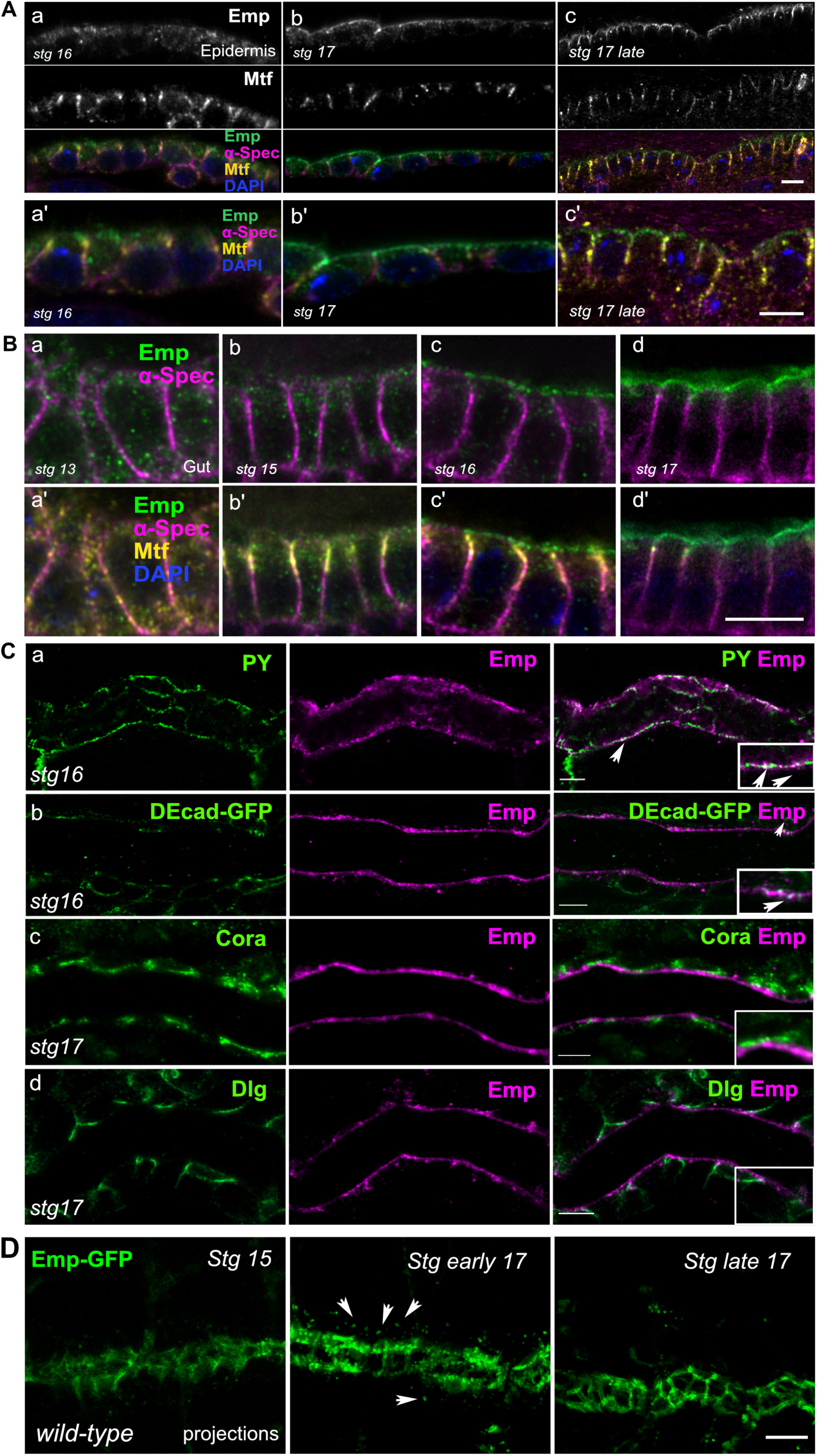
Emp localization. (A) Confocal images of embryonic epidermis stained for Emp, melanotransferrin (Mtf) and *α*-spectrin (*α*-Spec) showing the subcellular localization of Emp in stage 16 (a), stage 17 (b) and stage 17 late (c). (a’-c’) represents zoomed images from a – c. (B) Confocal images of embryonic gut stained for Emp, Mtf and *α*-Spec showing increasing apical accumulation of Emp in stage 13 (a-a’), stage 15 (b-b’), stage 16 (c-c’) and stage 17 (d-d’). (C) Confocal images of the tracheal DT, showing the relative localization of Emp and various cellular markers including PY (a), DE-cadherin (b), during stage 16, Cora (Coracle) (c) and Dlg (Disc Large) (d) during stage 17. Insets show zoomed cross section views of the DT of Emp co-localization with PY and DE-cadherin, as indicated by arrowheads. (D) Confocal frames of the dorsal trunk of living *wild-type* embryos expressing Emp-GFP (green). The protein gradually accumulates at apical junctions through intracellular trafficking. Arrowheads indicates the intercellular Emp. Scale bars, 5μm.

**Figure 3—figure supplement 1.**
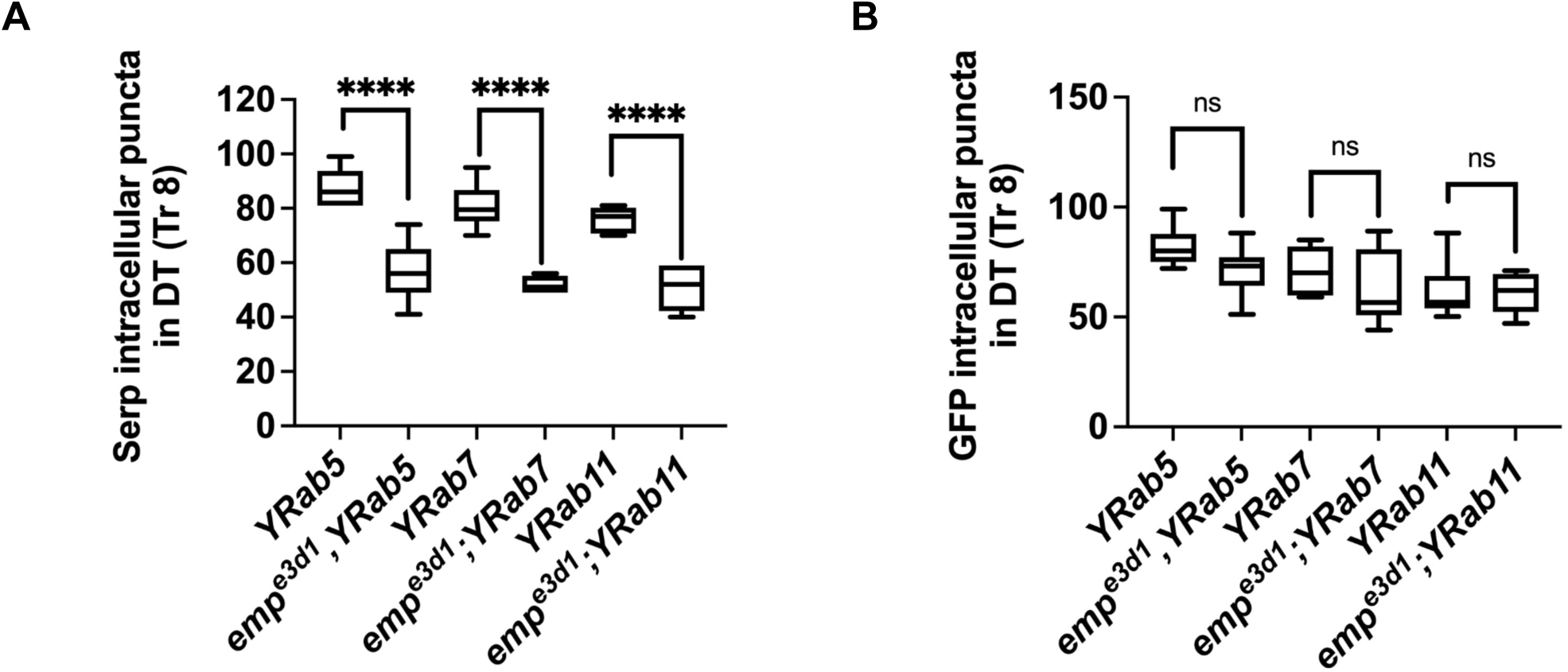
Quantification of Serp internalization. Boxplots of Serp (A) and GFP (B) intracellular puncta of tracheal cells in *YRab5*, *emp^e3d1^;YRab5*, *YRab7*, *emp^e3d1^;YRab7*, *YRab11* and *emp^e3d1^;YRab11* embryos at late stage 16. The boxplot (A-B) shows the median (horizontal line) and the data range from 25^th^ to 75^th^ percentile p>0,05 not significant (ns) and p <0,0001****(unpaired two tailed *t* tests).

**Figure 4—figure supplement 1.**
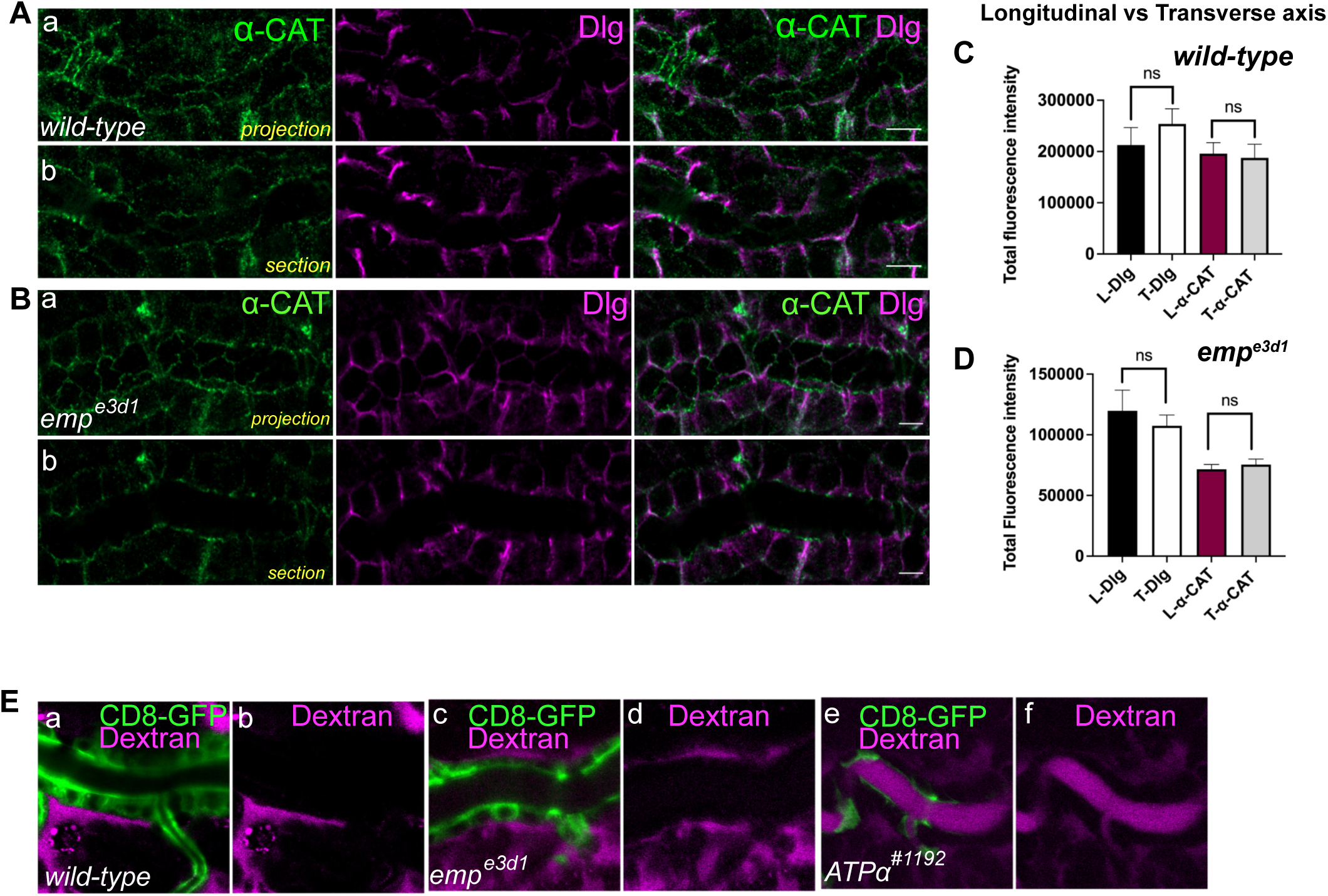
Epithelial cell integrity is not affected in *emp^e3d1^* mutants. (A) and (B) are confocal images of tracheal DT stained for *α*-CAT (green) and Dlg (magenta) respectively. Projections and sections of the tracheal tube in *wild-type* and *emp^e3d1^* embryos is shown. (C) and (D) plots showing the total fluorescence intensity of *α*-CAT and Dlg stainings in the Longitudinal (L-Dlg and L-*α*-CAT) and Transverse (T-Dlg and T-*α*-CAT) axis in *wild-type* and *emp^e3d1^* embryos respectively. (E) Selected confocal time-lapse images showing the tracheal clearance of Dextran-TR in living *wild-type*, *emp^e3d1^* and *ATPα* mutant embryos expressing the membrane marker *btl>CD8-GFP* during stage 16. Statistical significance shown in p-values p >0,05 not significant (ns) (Mann-Whitney test). Scale bars, 5 μm (A-B) and 10 μm (E). are intact in the mutants. Similarly, a dextran leakage assay comparing paracellular junction function in *wild-type*, *emp,* and *ATPα* mutant embryos showed that *emp* loss does not affect general SJ integrity (Figure 4-figure supplement 1E). Overall, these results suggest that Emp function regulates the apical membrane levels of Crb and DE-cad without majorly affecting junctional integrity or function. **Figure 4—source data 1** This zip archive contains the raw unedited western-blot shown in Figure 4D.

**Figure 6—figure supplement 1.**
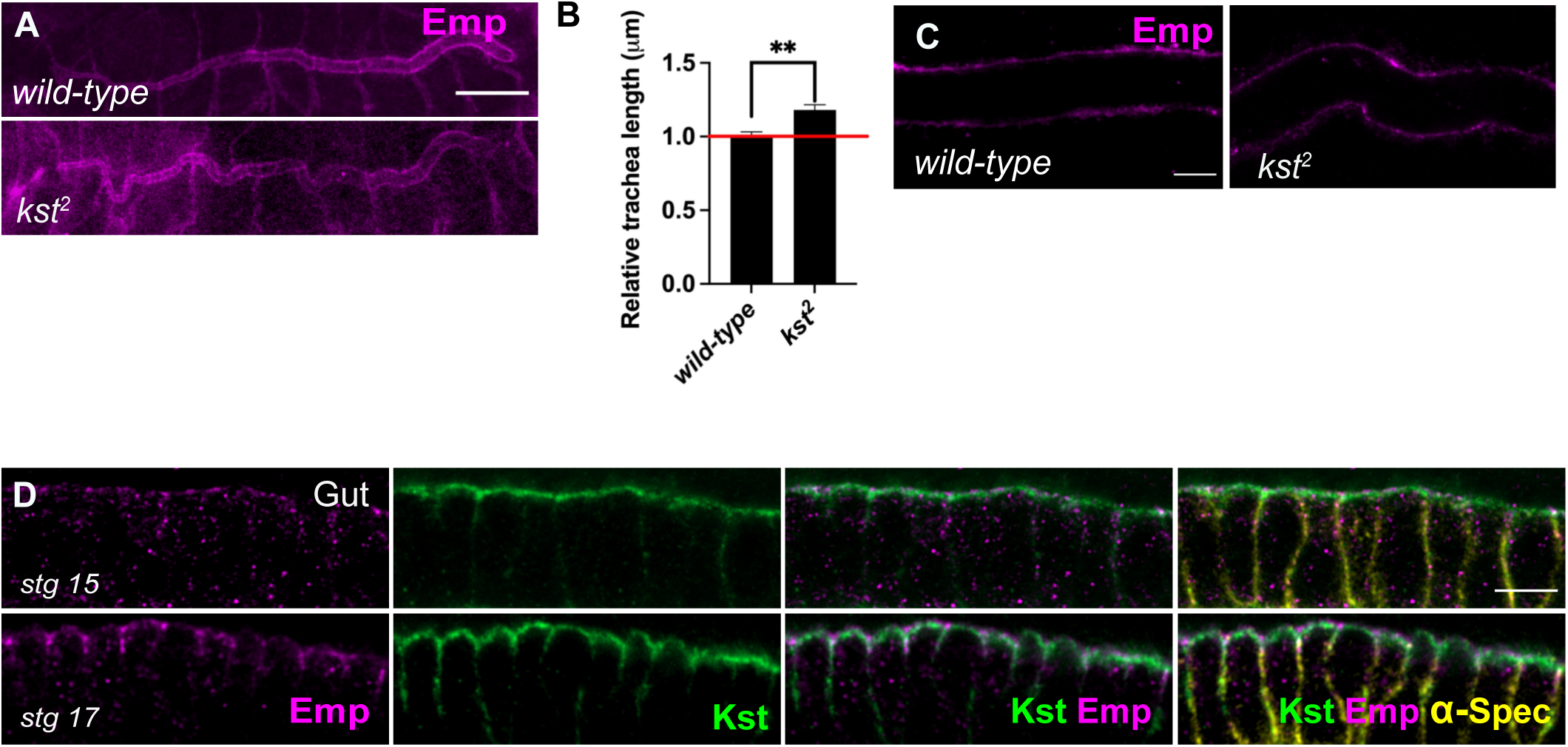
Tube elongation is defected in *kst^2^* mutants. (A) Confocal images showing the DT of *wild-type* and *kst^2^* embryos stained for Emp. (B) Bars plots are quantifications of the tracheal tube length in *wild-type* (n=6) and *kst^2^* (n=10) mutant embryos. (C) Confocal images showing the tracheal DT of *wild-type* and *kst^2^* embryos stained for Emp. (D) Confocal images showing the localization of Kst in the gut during stg 15 and stg 17, stained for Emp, GFP (Kst) and *α*-Spec. Error bars denotes s.e.m., p <0,005**(unpaired two tailed t tests). Scale bars, 50μm, (A) and 5µm (C, D). **Figure 6—source data 1** This zip archive contains the raw unedited western-blots shown in Figure 6E.

**Figure 7—figure supplement 1.**
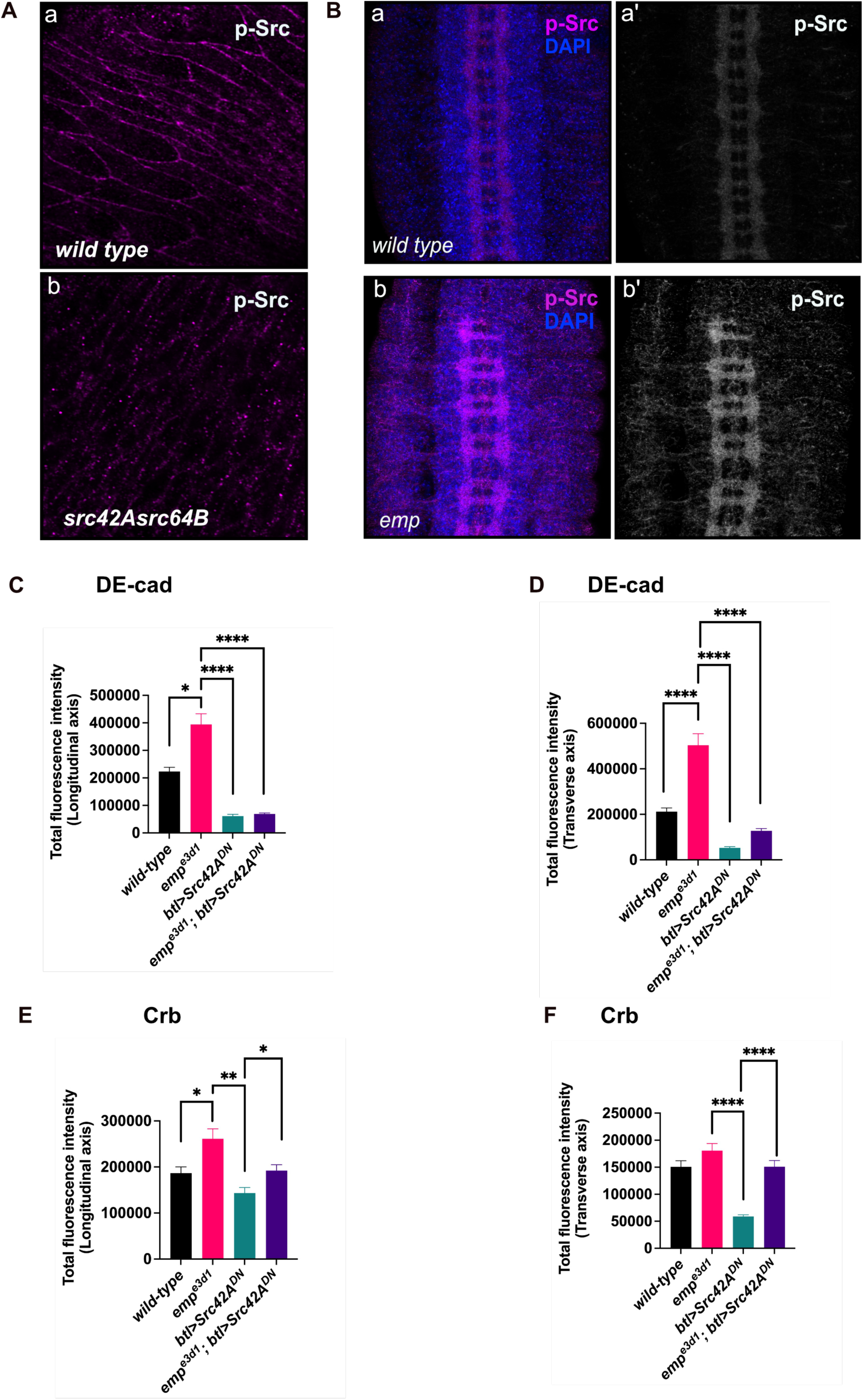
p-Src levels are increased in CNS of *emp^e3d1^* mutants. (A) Confocal images of embryonic epidermis stained for p-Src in *wild-type* (a) and *src42A^E1^;src64B* (b) embryos,showing the specificity of p-Src antibody. (B) Images of the ventral nerve cord in *wild-type* (a, a’) and *emp^e3d1^* embryos (b, b’) stained for p-Src and DAPI. (C) and (D) bar plots showing the total fluorescence intensity of DE-cad from DT in the longitudinal and transverse axis, respectively in *wild-type*, *emp^e3d1^*, *btl>src42ADN* and *emp^e3d1^;btl>src42A^DN^* embryos. (E) and (F) bars plots representing the total fluorescence intensity of Crb from DT in the longitudinal and transverse axis, respectively from the same embryos described in (C) (D). Error bars denotes s.e.m. wild-type, p < 0,05*, p < 0,005** and p < 0,0001****(Mann-Whitney tests). **Figure 7—source data 1** This zip archive contains the raw unprocessed western-blots shown in Figure 7B.

